# DynaKiller-Scan: Multimodal characterization of single-cell killing dynamics via combinatorial encoding of target lysis

**DOI:** 10.64898/2026.03.26.714656

**Authors:** Tongjin Wu, Lih Feng Cheow

## Abstract

Cytotoxic lymphocytes are crucial for eradicating infected or cancerous cells. To comprehensively characterize their killing capacity at single-cell level is challenging. Here we introduce DynaKiller-Scan, a high-throughput platform utilizing DNA barcodes to simultaneously interrogate the killing dynamics and molecular underpinnings of individual effector cells. DynaKiller-Scan reveals variable kinetics in serial killing between CD4^+^ and CD8^+^ CAR-T killers. The highly heterogeneous killing behaviors are associated with distinct molecular signatures orchestrating cell activation, secretion, migration, and metabolic fitness. While rapid killing involves cells with stronger type-I cytokine secretion, persistent killing is accompanied with more of type-II cytokine response. Furthermore, gene expression patterns closely linked to serial-killing were identified. DynaKiller-Scan could be a valuable tool for improving the development of precision immunotherapy.

## INTRODUCTION

Cytotoxic lymphocytes, mainly including cytotoxic T lymphocytes (CTLs) and natural killer (NK) cells, play a central role in immune surveillance against infected or malignant cells. These specialized immune cells can recognize and induce apoptosis in target cells through the release of cytolytic granules containing perforin and granzymes^1^. Today, a multitude of strategies have been exploited to harness their potential for better disease treatment and prophylaxis. These include adoptive cell therapy (ACT) using tumor-infiltrating lymphocytes (TILs), NK cells, or genetically re-directed chimeric antigen receptor (CAR)-T cells, CAR-NK cells, and T-cell receptor-engineered (TCR)-T cells^2–4^. Despite impressive advances, the broad application of cancer immunotherapies has been impeded by their highly variable response rates (e.g., low response and high relapse rate in solid tumors) and drug-induced severe toxicities (e.g., inflammation-mediated diseases)^2,5^.

The capabilities of cytotoxic lymphocytes have profound effects on the therapeutic outcomes in many immunotherapeutic approaches. One particularly fascinating behavior of cytotoxic lymphocytes is their ability to function as “serial killers”—a single lymphocyte can sequentially engage, kill, disengage, and eliminate multiple target cells^6–8^. This dynamic and efficient killing strategy was crucial for the immune system to regress solid tumors as observed in mouse models^8–10^, and correlated with clinical responses in CAR-T treated patients^11,12^. Thus, being able to systematically understand what makes an effector-cell capable of serial killing would offer new avenues for the development of optimal immunotherapeutic approaches.

Both CTLs and NK cells display substantial heterogeneity across multiple layers, including cellular diversity defined by phenotypic protein expression and molecular diversity defined by transcriptional profiles^13–15^, while only a subset of them were able to acquire serial killing activity^9–11,16^. Our understanding of these aspects remains fragmentary, and both the mechanisms governing serial killing and the degree of heterogeneity among individual cytotoxic lymphocytes are poorly understood. Therefore, progress in immuno-oncology research and therapeutic development requires the ability to robustly and cost-effectively identify and isolate target-specific killer cells. Moreover, there is a clear need for high-resolution, high-throughput, multi-modal approaches that can quantify cytotoxic activity at the single cell level, capturing the dynamics of killing events while directly linking functional phenotypes to underlying molecular features.

Conventional cytotoxicity assays, such as target-cell-centric chromium release assay (CRA)^17^ and lactate dehydrogenase (LDH) activity assay^18^, typically rely on population-level measurements and fail to capture the single-cell nature of killing events. Time-lapse imaging provides valuable insight into cytotoxic cell migration, search, and engagement with cognate targets at single-cell level^19–21^. More recently, the integration of nanowell grid array, microscopy imaging, micromanipulator, and single-cell RNA sequencing (scRNA-seq) makes it feasible to catch a glimpse of molecular mechanism-driven differential killing activity for individual CAR-T cells^9^. Nevertheless, this approach is inherently limited by low throughput and high cost in single-cell manipulation. Alternatively, we previously developed PAINTKiller (‘proximity affinity intracellular-transfer identification of killer cells’) assay^22^. In PAINTKiller, carboxyfluorescein succinimidyl (CFSE)-labelled target cells were co-cultured with effector cells that were surface-modified with CFSE-capturing antibodies, whereby individual killers were identified by their ability to capture the CFSE from lysed target cells. Despite its usefulness to identify individual killer cells, PAINTKiller lacks the capability to profile their killing dynamics.

In this study, we present a high-throughput method for simultaneously interrogating the killing dynamics and associated molecular underpinnings of individual effector cells (DynaKiller-Scan). DynaKiller-Scan starts from barcoding target-cell surface with fluorochromes or sequencing compatible oligonucleotides (Fig. 1a). Subsequently, co-culturing the modified target cells with effector cells enable trogocytosis of target labels by killer cells (Fig. 1b). The single-target killing or serial killing behavior, together with phenotypic protein expression of individual effectors, can be concurrently quantified by multiparameter flow cytometry or scRNA-seq workflow (Fig. 1c) for comprehensive interrogation of their cellular and molecular determinants (Fig. 1d, e). Using DynaKiller Scan, we found that although CD4^+^ and CD8^+^ CAR-T killers were transcriptionally distinct, they achieved comparable levels of multi-target killing activity. Notably, CAR-T subpopulations from both lineages exhibited pronounced heterogeneity in cytotoxic function, driven by a combination of shared and lineage specific genetic regulatory networks governing cell activation, polyfunctional secretion, migration, and metabolic fitness. Furthermore, DynaKiller-Scan uncovered distinct gene expression signatures associated with the progressive acquisition of serial killing behavior, enabling the stratification of highly potent CD8^+^ and CD4^+^ cytotoxic cells. By directly linking killing behavior with transcriptomic and phenotypic identities, this approach provides a powerful framework for dissecting cytotoxic cell heterogeneity and for guiding the rational optimization of cell-based immunotherapies.

**Figure 1.**
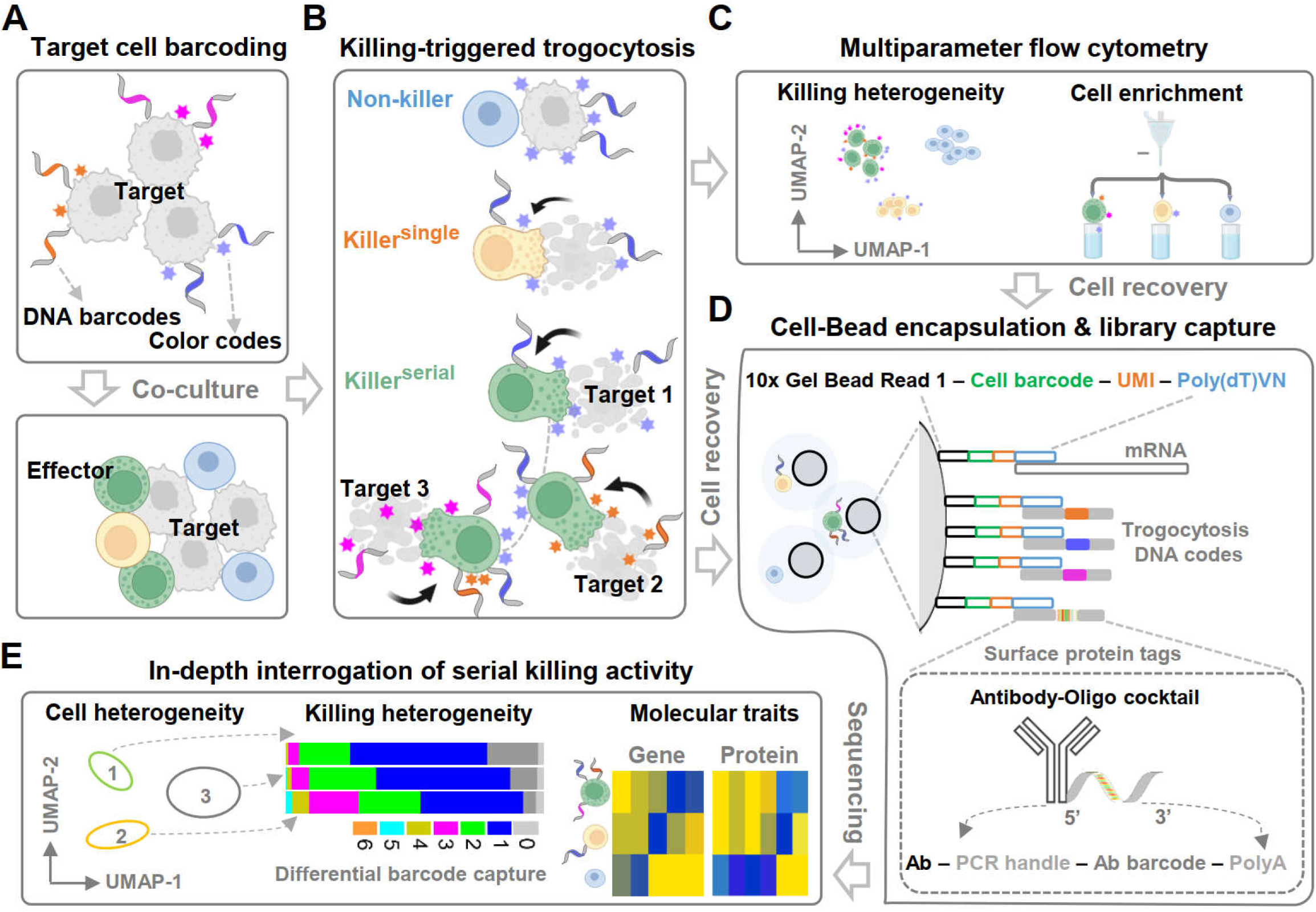
The principles of DynaKiller-Scan. **(A)** Target-cell barcoding with fluorochromes or oligonucleotides before co-cultured with effector cells. **(B)** Effector cells acquire target-cell labels via trogocytosis during killing. **(C)** Analysis of effector-cell killing dynamics by flow cytometry and enrichment of cytotoxic cells. **(D)** Multimodal profiling of single-cell killing behavior, cell surface marker and transcriptome. **(E)** Dissecting cytotoxic killing heterogeneity and the underlying cellular and molecular programs.

## RESULTS

### Selective transfer of target-cell labels to cytotoxic effector cells

Trogocytosis is a contact dependent process in which lymphocytes acquire membrane fragments from interacting cells during immune synapse formation. It occurs during lethal interactions between CTLs and cognate target cells^23,24^, as well as non-lethal cell-cell contacts^25^. Because cytotoxic engagements involve prolonged and stable synapse formation, they are expected to induce a substantially higher degree of trogocytosis than non-lethal interactions. Thus, in multicellular systems containing mixtures of killer and non-killer cells, trogocytosis has the potential to distinguish true cytotoxic activity from incidental cell-cell contact. However, the robustness of this distinction has not been systematically established.

To test this, we used engineered T cells (containing 20% CD19-directed CAR-T cells) as effectors and CD19-expresing NALM6 cells as targets. Given that trogocytosis involves the transfer of cell membrane, we adopt a general cell-surface engineering approach via whole-cell surface biotinylation followed by incubation with phycoerythrin-conjugated streptavidin (Strep-PE), which resulted in unique and stable Strep-PE labeling of NALM6 cell membrane (Fig. 2a,b). Both biotinylated and non-biotinylated NALM6 cells were efficiently killed by CAR-T cells, whereas no killing was observed with Non-CAR-T cells, indicating that biotinylation does not affect CAR-T–mediated cytotoxicity (Fig. 2c,d). If trogocytosis is associated with CAR-T cell cytotoxic activity, we would expect an increased fraction of CAR-T cells to acquire Strep-PE. Consistent with this, using the non-CAR-T capture background (∼0.5%) to define the gate, a substantial proportion of CAR-T cells were Strep-PE positive (Fig. 2e,f).

**Figure 2.**
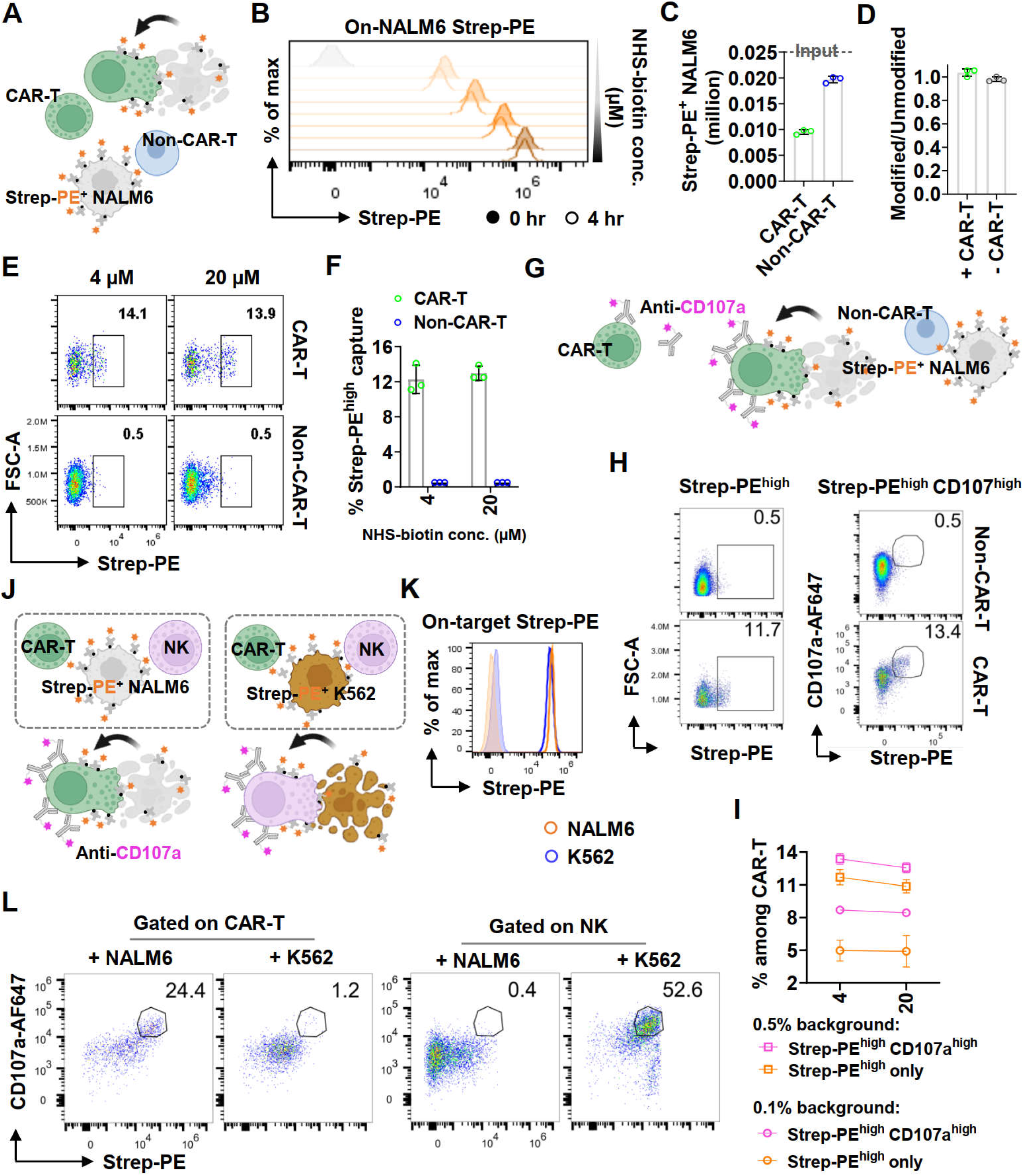
Selective transfer of target-cell labels to cytotoxic effector cells. **(A)** Co-culture of CAR-T, Non-CAR-T, and Strep-PE-modified NALM6 cells (1:4:1). **(B)** Target cells were modified with different concentrations of NHS-biotin (500, 100, 20, and 4 µM). Strep-PE label on target cell before and after co-cultured with CAR-T cells. **(C)** CAR-T cells were able to specifically kill Strep-PE modified NALM6 cells. **(D)** Unmodified and Strep-PE modified NALM6 cells were mixed at equal proportion and co-cultured with CAR-T or Non-CAR-T cells. Similar killing efficiency of unmodified and modified NALM6 cells by CAR-T cells was observed. **(E-F)** Shown were the gating to identify trogocytosis of Strep-PE by CAR-T cells based on background capture of Non-CAR-T counterpart **(E),** and summary of three independent experiments **(F). (G)** Improving the assay specificity by co-labeling effector cells with CD107a antibodies. **(H-1)** Gating strategy to identify CD10?high Strep-PEhigh killer cells **(H),** and inclusion of CD107a as a second marker improved the sensitivity and specificity of detecting killer cells (I). **(J)** Experimental schematic of co-culturing labeled target cells with different effector cells (CAR-T and NK cells). **(K)** Universal target labeling scheme was applicable across different cell types. **(L)** Labels from NALM6 cells were transferred exclusively to CAR-T cells, whereas labels from K562 cells were transferred exclusively to NK cells, consistent with trogocytosis occurring during cognate cytotoxic interactions.

Cytotoxic cell degranulation, accompanied by the expression of CD107a, is a key process during functional killing^26^. Co-staining effector cells with anti-CD107a revealed that Strep-PE^high^ capture overlapped with upregulated CD107a expression on CAR-T cells (Fig. 2g,h; Supplementary Fig. 2), confirming that the Strep-PE^high^ effectors cells were active cytotoxic CAR-T cells engaged in killing. Notably, using the same background threshold, this dual-marker scheme reliably identified a larger and more consistent proportion of killer cells while minimizing signals from potential non-cytotoxic interactions (Fig. 2i). Furthermore, we mixed CAR-T and NK cells together with labelled NALM6 (CAR-T target) or K562 (NK target) (Fig. 2j,k), and demonstrated that this dual-marker approach also allowed us to robustly identify matched killer-target from a heterotypic CAR-T/NK mixture (Fig. 2l), opening the door for future work in identifying functional specificity across multi-effectors and multi-targets.

### Multicolor fluorescence encoding of serial target-cell killing

A subset of cytotoxic lymphocytes is capable of serial killing^9,11^. In our assay, each killer cell acquires membrane labels from the target cells it kills. We hypothesized that serial killers—cells that kill multiple distinct targets—could be identified by the presence of multiple fluorophore-conjugated labels acquired via trogocytosis (Fig. 3a). To test this, different populations of biotinylated NALM6 cells were labeled with distinct streptavidin-fluorophore conjugates (Strep-Fluo; e.g., PE, BV421, and BV786), and co-cultured with CAR-T cells in the presence of anti-CD107a. These labeled targets remained well-separated, with minimal cross-talk even at high cell densities, indicating the stability of labeling (Fig. 3b; Supplementary Fig. 2). Upon analysis, distinct CAR-T cell subpopulations that were positive for PE, BV421, or BV786 emerged, corresponding to the killing of the respective target populations (Fig. 3c). Non–CAR-T cells showed negligible CD107a expression and minimal dye acquisition, underscoring the specificity of our assay (Fig. 3c).

**Figure 3.**
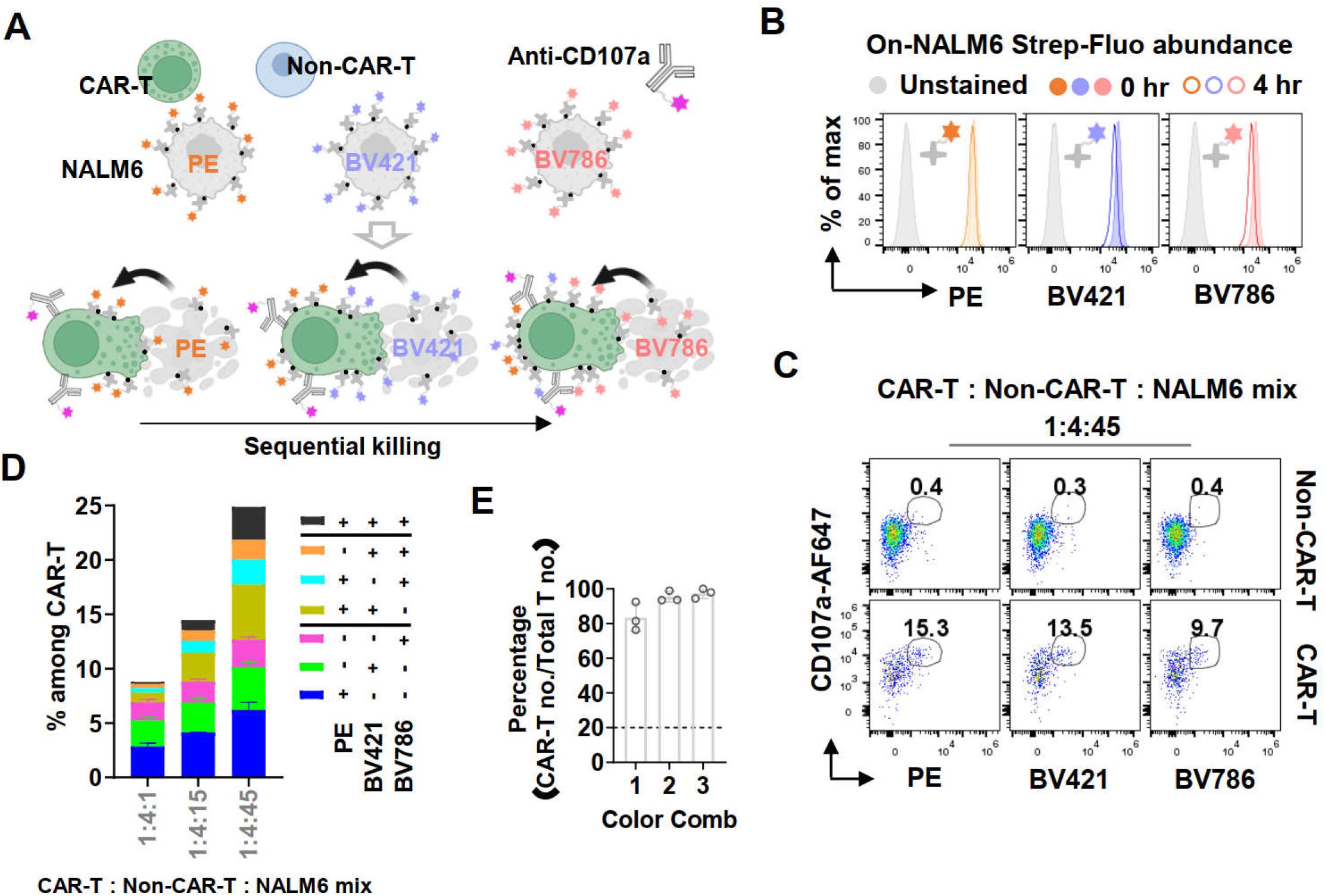
Multicolor fluorescence encoding of serial target-cell killing. **(A)** Shown was co-culture of CAR-T, Non-CAR-T, and differentially labelled NALM6 (equally mixed Strep-PE, Strep-BV421, and Strep-BV786 modified cells) at high E:T ratio (1:4:1), enabling the identification of effector cells that had killed multiple targets by accumulation of multiple colors. **(B)** High stability of on-target Strep-Fluo before and after co-culture. **(C-D)** Co-culture of CAR-T, Non-CAR-T, and differentially labelled NALM6 at low E:T ratio (1:4:45) in the presence of anti-CD107a-AF647. Gating to identify the acquisition of each Strep-Fluo by CAR-T cells **(C)** and summary of killing events across different E:T ratio **(D).** (E) Acquisition of single and multiple target-cell fluorophores was highly restricted to CAR-T cells.

We quantified the number of CAR-T cells with single and multiple fluorophore combinations (Fig. 3d). At high E:T ratios, where targets were limiting, most killer cells acquired only one color. In contrast, at low E:T ratios (target-rich conditions), we observed a significant increase in two or more fluorophores-labeled CAR-T cells, consistent with enhanced serial killing (Fig. 3d). Subsequent experiments were therefore conducted at low E:T ratios to avoid target limitation and better capture serial killing dynamics. We quantified the proportion of dye-labeled CAR-T in dye-labeled total T cells for each fluorophore combination and found high values across the board (Fig. 3e), indicating the feasibility and specificity of multiplex trogocytosis-based serial killing identification.

### DNA barcode encoding of serial target-cell killing

While the multiplexed fluorophore-based approach is robust, it is ultimately limited by the number of available dyes that do not overlap spectrally. This restricts the number of target populations that can be simultaneously tracked and constrains the resolution of the assay. Furthermore, flow cytometry analysis only enables correlation of cytotoxic activity with surface phenotypes, limiting insight into underlying molecular programs.

To overcome these limitations, we developed a DNA-based labeling strategy. In this system, biotinylated NALM6 target cells were labeled with Strep-PE, followed by incubation with a biotinylated DNA oligo (Supplementary Fig. 3). We then introduced a complementary Cy5 fluorophore-conjugated DNA probe designed to hybridize with the cell-surface DNA barcode (Fig. 4a, illustration). Flow cytometric analysis revealed co-staining and strongly correlated fluorescence intensity of PE and Cy5 on target cells (Fig. 4a), confirming that the amount of DNA on the cell surface is tunable and controllable through the degree of biotin-streptavidin loading.

**Figure 4.**
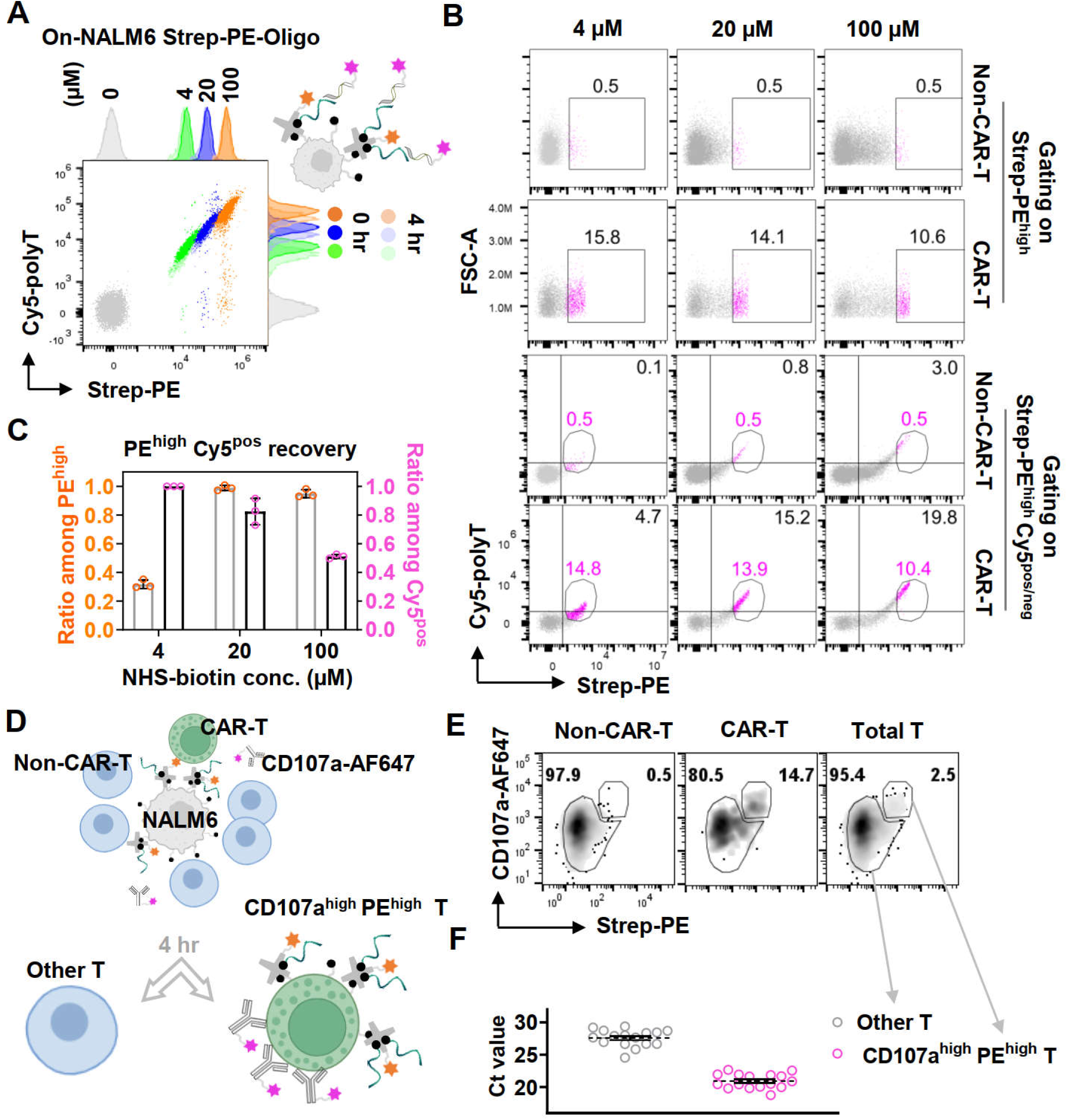
DNA barcode encoding of serial target-cell killing. **(A)** NALM6 cells were modified with different concentrations of NHS-biotin before labeling with Strep-PE to support the attachment of biotin-conjugated DNA barcodes, followed by hybridization with Cy5-polyT probe. Both Strep-PE and DNA barcodes were stably anchored on cell surface over 4 hours. **(B)** Gating to identify the trogocytosis of Strep-PE or Strep-PE-Oligo complex by CAR-T cells. **(C)** The percentage of Strep-PEhigh Cy5P^05^ CAR-T cells among total Strep-PEhigh or Cy5P^05^ CAR-T cells. **(D-F)** Co-culture of CAR-T, Non-CAR-T, and DNA-labelled NALM6 (1:4:1) in the presence of anti-CD107a-AF647 **(D),** gating to sort single T cell with high or low CD107a and Strep-PE co-staining after 4 hr (E), and q-PCR to quantitate the relative DNA barcode abundance as described in Methods **(F).**

Next, we performed a killing assay by incubating the DNA-labeled NALM6 cells with engineered effector T cells. As expected, we observed a distinct population of CAR-T cells that was positive for both PE and Cy5 signal, indicating that DNA barcodes on the target cell surface were transferred along with membrane proteins during trogocytosis (Fig. 4b). While both PE (protein) and Cy5 (DNA) signals were reliably detectable, we noted potential differences in detection efficiency between the two. To optimize target cell labeling, we compared the proportion of double-positive (PE^high^ Cy5^+^) cells among PE^high^ and Cy5^+^ populations across the different biotinylation condition (Fig. 4c,d). We found that an intermediate labeling concentration (20 μM) provided the best balance and sensitivity, maximizing the overlap between protein and DNA detection.

To further validate the detected Cy5 signal was equivalent to the captured target barcodes, effector T cells after co-culture were sorted as single cells into PE^high^ CD107a^high^ (confirmed killers) and the PE^low^ CD107a^low^ counterpart (non-killers) (Fig. 4e,f). The abundance of captured target DNA barcodes was detected with qPCR using matched primer sets. The clear separation in qPCR cycle threshold (Ct) values between the two group further confirms that the DNA barcode is specifically and reliably transferred to killer effector cells during target engagement (Fig. 4g).

Together, these results demonstrate that our DNA-trogocytosis platform provides a robust, specific, and molecularly validated approach for tracking effector–target interactions and identifying the precise targets killed by individual cytotoxic lymphocytes. As there are much more DNA barcodes compared to distinct fluorophores, this strategy would significantly increase multiplexing capacity, allowing for the simultaneous tracking of many distinct target cell populations far beyond what is feasible with fluorophores. Combined with scRNA-seq, this also sets the foundation for integrating functional readouts with transcriptomic profiling at single-cell resolution.

### DynaKiller-Scan characterizes lineage-specific cytotoxic behavior and molecular traits in CAR-T cells

Having validated the specificity and robustness of DNA barcode transfer via trogocytosis, we next sought to link the cytotoxic function to molecular profiles at single-cell resolution. To achieve this, we integrated our DNA-trogocytosis platform with CITE-seq^27^ using the 10x Genomics Chromium platform. This allowed us to capture full transcriptomes alongside transferred DNA barcode sequences and membrane phenotypic protein-targeted antibody DNA tags within individual effector cells (Fig. 1d).

In this experiment (Fig. 5a), NALM6 target cells were first biotinylated and divided into six equal populations, each labeled with a unique Strep-PE-DNA oligo barcode. These membrane DNA oligos served dual functions: acting as molecular identifiers of the specific target populations and enabling detection of trogocytosis events through single-cell sequencing. The labeled NALM6 cells were pooled and co-incubated with purified CAR-T or non-CAR-T cells in a standard killing assay (1 and 4 hr) in the presence of anti-CD107a. At the end of the assay, as a large proportion of CAR-T cells did not engage in killing, we focused on the most functionally relevant subpopulation by using FACS for CD107a^high^ PE^high^ CAR-T cells—those that had acquired both PE signal, indicating membrane uptake from target cells, and CD107a expression, indicating active degranulation (Supplementary Fig. 4a,b). These cells represented the killer effector population. In parallel, we also sorted a control group of the CD107a^low^ PE^low^ CAR-T cells, which did not exhibit signs of cytotoxic engagement and could serve as a baseline reference for transcriptomic comparison. These cell samples were individually barcoded with TotalSeq-A hashtags. The cells were pooled and stained with a cocktail of human immuno-profiling TotalSeq-A antibodies for high-throughput single-cell multiomics sequencing.

**Figure 5.**
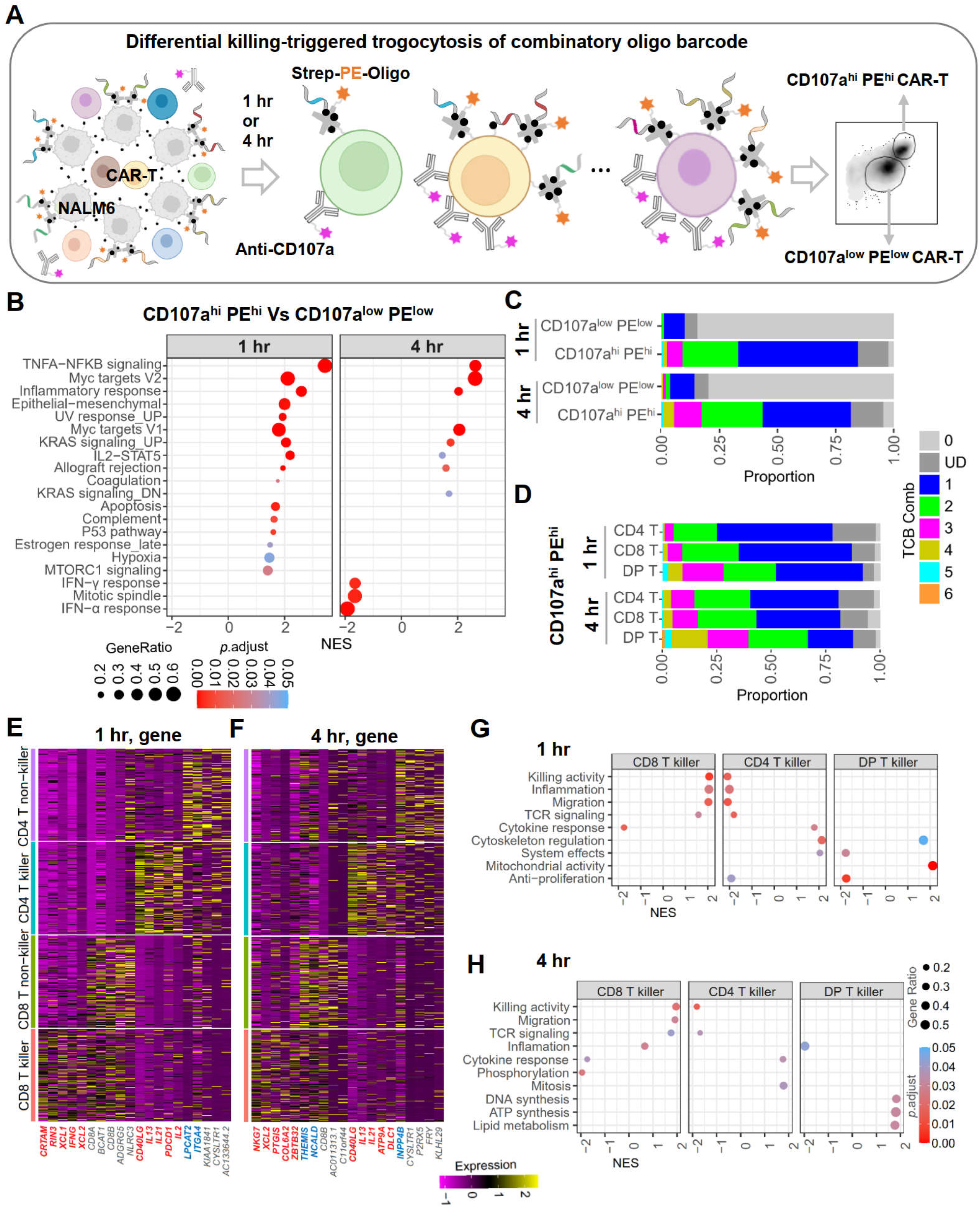
DynaKiller-Scan reveals lineage-specific cytotoxic behavior and molecular traits in CAR-T cells. **(A)** Co-culture of CAR-T, Non-CAR-T, and DNA-barcoded NALM6 at low E:T ratio (1:4:45) in the presence of anti-CD107a-AF647, and gating to sort CD107ahigh Strep-PEhigh and CD107a^10^w Strep-PE^10^w CAR-T cells after 1 or 4 hr (also refer to Supplementary Fig. 4a,b). **(B)** Hallmark gene sets enriched in isolated CAR-T populations. **(C-D)** Distribution of combinatorial target-cell barcode (TCB) capture in sorted CAR-T cells (C) and lineage-annotated subsets (D), grouped by the number of barcode captures (1-6) or classified as undetermined (UD). **(E-F)** Representative genes differentially expressed by cos+ CAR-T killers, cos+ CAR-T non-killers, CD4+ CAR-T killers, and CD4+ CAR-T non-killers. Genes uniquely upregulated by cos+ or CD4+ CAR-T killers were colored in red while those downregulated were in blue. **(G-H)** GO biological processes enriched by cos+ CAR-T killers, CD4+ CAR-T non-killers, and DP CAR-T killers.

To begin our analysis, we demultiplexed the sorted CD107a^high^ PE^high^ and CD107a^low^ PE^low^ CAR-T cells for each timepoint after sequencing (Supplementary Fig. 4c,d), and compared their transcriptomes. The CD107a^high^ PE^high^ CAR-T cells were highly enriched for genes linked to cell activation (e.g., TNF-α/NF-κB signaling, *MYC* targets, inflammatory response, and *KRAS* activation signaling), while CD107a^low^ PE^low^ CAR-T cells upregulated genes favoring immune-regulatory or inhibitory function (e.g., *NLRC3* and *MYO1F*) (Fig. 5b and Supplementary Fig. 5). Intriguingly, CD107a^high^ PE^high^ CAR-T cells exhibited an attenuated interferon response over time (Fig. 5b), potentially reflecting a mechanism of enhanced tolerance to interferon stimulation to preserve their long-term effector function^28–30^. Together, these findings confirm that the functionally defined killer population, according to trogocytosis and degranulation level, was transcriptionally programmed for robust effector activity, providing a foundation for further dissection of heterogeneity within the killer pool.

To quantify the serial killing behavior of individual T cells, we carefully analyzed the trogocytosis of target-cell barcodes (TCB) and corresponding transcriptomic change (Supplementary Fig. 6,7; refer to Methods). Each unique barcode corresponds to a distinct target cell population. Thus, the number of TCB acquired by a given effector cell reflects the number of distinct targets it had killed. As expected, the CD107a^low^ PE^low^ CAR-T (non-killer bystanders) showed almost exclusively negative barcode capture, consistent with a lack of cytotoxic engagement (Supplementary Fig. 6b). In contrast, the CD107a^high^ PE^high^ CAR-T population showed widespread barcode acquisition, confirming successful trogocytosis and functional target recognition (Fig. 5c). Most of these killer cells captured one barcode, reflecting engagement with a single target. However, a substantial fraction captured two barcodes, and a smaller but clear subsets captured three or four barcodes. The proportion of cells that captured multiple TCB increased over time, indicating multi-target killing behavior. Barcode capture of five or six was rare, but detectable, representing highly potent serial killers.

We further stratified the CAR-T cells into CD4^+^ T, CD8^+^ T, CD4^+^ CD8^+^ double-positive T (“DP T”), and CD4^−^ CD8^−^ double-negative T (“DN T”) according to their expression of lineage-specific genes (*CD4*, *CD8A*, and *CD8B*) and cell surface proteins (CD4 and CD8a) (Supplementary Fig. 8). The identified T-cell types were transcriptionally distinct (Supplementary Fig. 9a-c), and computational analyses indicated a low frequency of cell doublets (Supplementary Fig. 9d,e). There was an increase in CD8^+^ to CD4^+^ CAR-T ratio among the CD107a^high^ PE^high^ CAR-T population (killer cells) compared to its CD107a^low^ PE^low^ counterpart (non-killer cells) (Supplementary Fig. 10a,b, histogram plots), suggesting CD8^+^ CAR-T were the dominant killer cells in our system. Unexpectedly, CD8^+^ and CD4^+^ CAR-T manifested a very similar pattern of target barcode capture over time (Fig. 5d and Supplementary Fig. 10), suggesting qualitatively comparable serial killing potential of generated CD4^+^ CAR-T over CD8^+^ CAR-T cells. Interestingly, although DP CAR-T cells constituted a rare population, they became enriched within the CD107a^high^ PE^high^ group during late-stage co-culture (Supplementary Fig. 10b, histogram) and comprised a substantial fraction of cells classified as serial killers, suggesting enhanced cytotoxic potency (Fig. 5d). This DP T cells have also been observed in other studies, especially in harsh tumor microenvironment^31,32^.

Genes that differentiated CD8^+^ killers from CD4^+^ killers would help elucidate how therapeutic T cells of different lineages may function together to cure diseases^33–35^. We found that TCB capture-defined CD8⁺ killers were distinguishable from CD4^+^ killers in the expression of core gene signatures over time (Fig. 5e,f and Supplementary Fig. 11). GSEA showed that CD8⁺ killers were enriched for biological processes such as cell cytotoxicity, migration, inflammation, and TCR signaling, whereas CD4⁺ killers had increased cytokine-mediated signaling, cytoskeleton remodeling, and broader systems-level immune functions. Instead, DP killers had upregulated genes relating to mitochondrial activity and lipid metabolism (Fig. 5g,h and Supplementary Fig. 12). These gene modules suggest differing functional programs aligned with their respective T cell identities, and pointing to a more immunoregulatory and supportive role for CD4⁺ killer cells, distinct from the direct cytolytic focus of their CD8⁺ counterparts. Together, our results showed that although CD8^+^ and CD4^+^ CAR-T cells had comparable levels of multi-target killing activity, they were driven by distinct molecular programs and manifested by different kinetics.

### DynaKiller-Scan reveals diverse killing dynamics among CAR-T subpopulations

To further dissect the cellular and molecular heterogeneity underlying serial killing capacity, we performed UMAP clustering for CD4⁺ and CD8⁺ CAR-T cells based on their transcriptional profiles. This analysis revealed 10 distinct clusters within the CD8⁺ compartment and 9 clusters among CD4⁺ cells, reflecting a diverse range of transcriptional states even within functionally defined killer subsets (Fig. 6a,b, dot plots, and Supplementary Fig. 13). By quantifying the average number of DNA barcodes captured per cell in each cluster, we observed substantial variation in cytotoxic behavior across clusters (Fig. 6a,b, histogram plots). Some clusters, such as CD8 cluster 1 and 0, or CD4 cluster 4 and 1, showed an average of 1.8 barcodes per cell, whereas other clusters, like CD8 cluster 5 and CD4 cluster 6, were dominated by cells with low or no barcode capture (Fig. 6a,b, line charts). These findings suggest that cytotoxic potential is not evenly distributed but rather enriched within specific transcriptional states.

**Figure 6.**
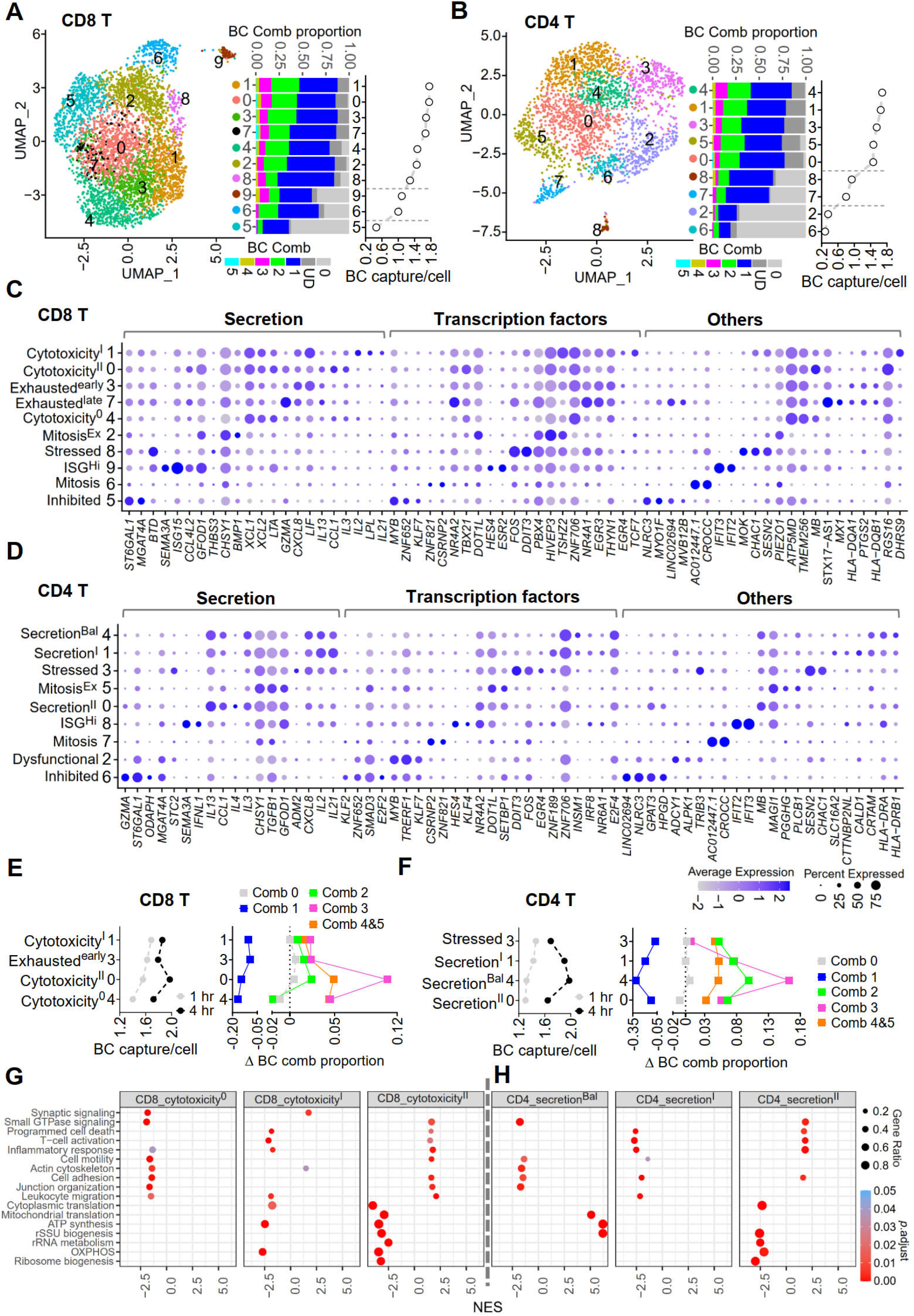
DynaKiller-Scan reveals diverse killing dynamics among CAR-T subpopulations (A-B) Unsupervised coa+ and co4+ CAR-T cell clustering based on global transcriptional profile (UMAP plots), and combinatorial capture of TCB for each cluster (histogram plots). The clusters were ranked by the average number of captured TCB from high to low (line plots). **(C-D)** Cell cluster annotation according to top differentially expressed genes that were grouped into secreted proteins, transcription factors, and intracellular/membrane-bound proteins. **(E-F)** Time-dependent change of average TCB capture for coa+ or co4+ CAR-T cell clusters that were characterized by high cytotoxicicity and cytokine-secretion gene signatures **(G-H)** GO biological processes enriched in coa+ and co4+ CAR-T clusters that were most enriched with serial killers.

To annotate the refined CAR-T cell clusters by their function, differentially expressed genes were identified and categorized into functional secretion, transcription factors, and intracellular/surface proteins (Fig. 6c,d; refer to Methods). In addition, we performed Jaccard similarity analysis of the gene expression between the identified CD8^+^ and CD4^+^ clusters to help find common cell annotation (Supplementary Fig. 14).

As depicted in Fig. 6c, CD8^+^ CAR-T cells were annotated as **(1) Cytotoxicity^I^**, **Cytotoxicity^II^**, and **Cytotoxicity**^0^ cells that shared common effector secretions (e.g., *XCL1*, *XCL2*, and *LTA*). Cytotoxicity^I^ was characterized by higher gene expression for type-I cytokines (e.g., *IL2* and *IL21*)^36,37^, regulator cytokine (*LIF*), TFs of early T cell development and differentiation (*TSHZ2* and *TCF7*); Conversely, Cytotoxicity^II^ shifted to express more of the type-II cytokines (e.g., *IL13* and *IL3*)^38,39^ but with reduced *LIF*, *TSHZ2*, and *TCF7*. Cytotoxicity^0^ had globally reduced functional secretion. **(2) Exhausted^early^** and **Exhausted^late^** cells were marked by weakened cytokine secretion. Compared to Exhausted^early^ cells, Exhausted^late^ cells highly expressed *GZMA* and T cell activation negative regulators (e.g., *EGR3*, *NR4A1*, and *NR4A2*). **(3) Stressed** cells overexpressed stress-responding genes (e.g., *FOS*, *DDIT3*, *CHAC1*, and *SESN2)* that were associated with cell senescence or functional suppression. **(4) ISG^Hi^** cells were enriched of interferon-stimulated genes (e.g., *ISG15*, *IFIT3*, and *IFIT2*), and TFs (e.g., *HES4* and *ESR2*) potentially leading to T-cell over-activation^40,41^. **(5) Inhibited** cells were marked by high abundance of *NLRC3*, a known negative regulator of T-cell function^42^. **(6) Mitosis** or mitosis-exiting (**Mitosis^Ex^**) cells had largely reduced transcript amount (Supplementary Fig. 15), corresponding to global transcription silencing during cell division^43^. Mitosis cells were also distinguishable by highly upregulated *CROCC* (encoding rootletin), a component required for centrosome cohesion.

Likewise, as shown in Fig. 6d, CD4^+^ CAR-T cells were annotated as **(1) Secretion^I^** cells highly enriched for type-I cytokine (e.g., *IL2* and *IL21*)^36,37^, **Secretion^II^** cells enriched for type-II cytokine (e.g., *IL13*, *IL3*, and *IL4*)^38,39^, and **Secretion^Bal^** cells with a balanced type-I/II secretion. Secretion^II^ cells, like CD8^+^ Cytotoxicity^II^ cells, exhibited high *MB* gene expression. **(2)** CD4^+^ **Stressed** cells, like its CD8^+^ counterpart, generally reduced polyfunctional cytokine secretion, but upregulated stress-responding genes. **(3)** CD4^+^ **ISG^Hi^** cells resembled CD8^+^ ISG^Hi^. **(4)** Compared to CD8^+^ Inhibited cells, there were two similar CD4^+^ counterparts. One was **Inhibited** cluster highly expressed *GZMA* and some known negative regulators (e.g., *NLRC3*, *C15orf53*, and *HPGD*) of T-cell effector function. The other was **Dysfunctional** cluster manifested by upregulated *MYB* and *TRERF1* but reduced *NLRC3, C15orf53, and HPGD*. **(5) Mitosis** and mitotic exit cells (**Mitosis^Ex^**).

By integrating detailed cell annotation with measures of cytotoxic potency, we resolved T cell subpopulations with heterogeneous functional killing behaviors (Fig. 6a,b, line charts). In general, **Inhibited** and **Dysfunctional** clusters performed poorly in target-cell elimination, followed by **Mitosis** and **ISG^Hi^** clusters. Highly secretory clusters such as CD8^+^ **Cytotoxicity^I^**, **Cytotoxicity^II^**, **Cytotoxicity**^0^, and CD4^+^ **Secretion^Bal^, Secretion^I^, Secretion^II^** exhibited robust serial killing behavior, and were marked by gene expression signatures related to strong bioenergetic production and ribosomal activities (Supplementary Fig. 16). CD8^+^ **Exhausted** cells and CD4^+^ **Stressed** cells showed compromised killing effect compared to their highly secretory states respectively.

Furthermore, the ability of cytotoxic CD8⁺ and CD4⁺ T cells to sustain serial killing over time was strongly dependent on their cellular state (Fig. 6e,f and Supplementary Fig. 17). At the early stage (1 h), cytotoxic cells exhibiting type-I cytokine secretion programs (**Cytotoxicity^I^** and **Secretion^I^**) displayed the most rapid killing kinetics, whereas at the later stage (4 h), cytotoxic cells with upregulated type-II cytokine secretion programs (**Cytotoxicity^II^** and **Secretion^Bal^**) unexpectedly showed the highest serial killing potency. These temporal differences in killing behavior were mirrored by distinct molecular programs. In CD8⁺ T cells, **Cytotoxicity^I^** killers exhibited the strongest immunological synapse activity, consistent with their rapid early target-cell elimination. In contrast, **Cytotoxicity^II^** killers showed enhanced T-cell activation, adhesion, junction formation, cellular organization, and migratory capacity, features that are likely to facilitate efficient disengagement and re-engagement with multiple targets, thereby supporting sustained serial killing at later time points. Notably, this state was accompanied by reduced oxidative phosphorylation, ATP synthesis, and ribosome biogenesis, suggesting a metabolic adaptation that may favor prolonged functionality over acute effector output (Fig. 6g). Among CD4⁺ cytotoxic T cells, **Secretion^Bal^** cells displayed reduced motility, adhesion, actin cytoskeleton remodeling, and junction organization compared with **Secretion^I^** and **Secretion^II^** cells, indicating slower translocation dynamics. However, these cells were enriched for genes involved in ATP production, mitochondrial activity, and ribosome biogenesis, which may support prolonged metabolic fitness and compensate for reduced migratory efficiency, thereby enabling sustained cytotoxic activity over time (Fig. 6h). Together, these findings suggest that early cytotoxic efficiency could be driven by strong synaptic engagement, whereas long-term serial killing capacity may depend on a coordinated balance of migration, activation state, and metabolic adaptation, highlighting distinct molecular strategies underlying acute versus sustained CTL-mediated target-cell killing.

### Dynamic gene expression underlies CAR-T serial killing capacity

While a proportion of cells from each of the above identified T-cell clusters were endowed with serial target-killing capacity (Fig. 6a,b and Supplementary Fig. 18), we were wondering if there exist unique or shared molecular programs that may underpin the serial killing state for CD8^+^ or CD4^+^ CAR-T cells.

We thus calculated the relative change of individual gene along with sequential target-cell killing, and kept only those passing significant test (Fig. 7a). We found that, with the increase of target barcode capture, there were a few main different patterns of gene expression dynamics, including immediate upregulation/downregulation (“Imm_up”/“Imm_down”), delayed upregulation/downregulation (“Del_up”/“Del_down”), transient upregulation/downregulation (“Tra_up”/“Tra_down”), and fluctuating state (“Fluct”) (Fig. 7b). As depicted, most significantly changed genes were identified as immediate upregulation, immediate downregulation, or delayed upregulation. Unexpectedly, while most of the genes (1751) were shared between CD8^+^ and CD4^+^ killers, most of them were expressed in different dynamic patterns (Fig. 7b). For instance, *CCL3*, *CCL4*, and *NR4A3* showed gradual upregulation before achieving plateau during the sequential target killing (“Gradual” up) in CD8^+^ killers, but with rapid increase at only initial stage of target encountering (“Rapid” up) in CD4^+^ killers. On the contrary, *RNF213*, *SOS1*, and *BCL2* downregulated at an earlier stage (“Rapid” down) for CD8^+^ killers when compared to that for CD4^+^ killers (Fig. 7c,d). Interestingly, some genes like *TRAF3*, *ARID5B*, *RPS19BP1*, *CD48*, and *TMSBX4* showed even an opposite expression pattern in CD8^+^ and CD4^+^ killers (Supplementary Fig. 19). These observations highlight pronounced transcriptional plasticity during sequential target engagement, and reveal that CD8⁺ and CD4⁺ CAR-T cells achieve serial killing through divergent temporal expression strategies, even when leveraging the same molecular components.

**Figure 7.**
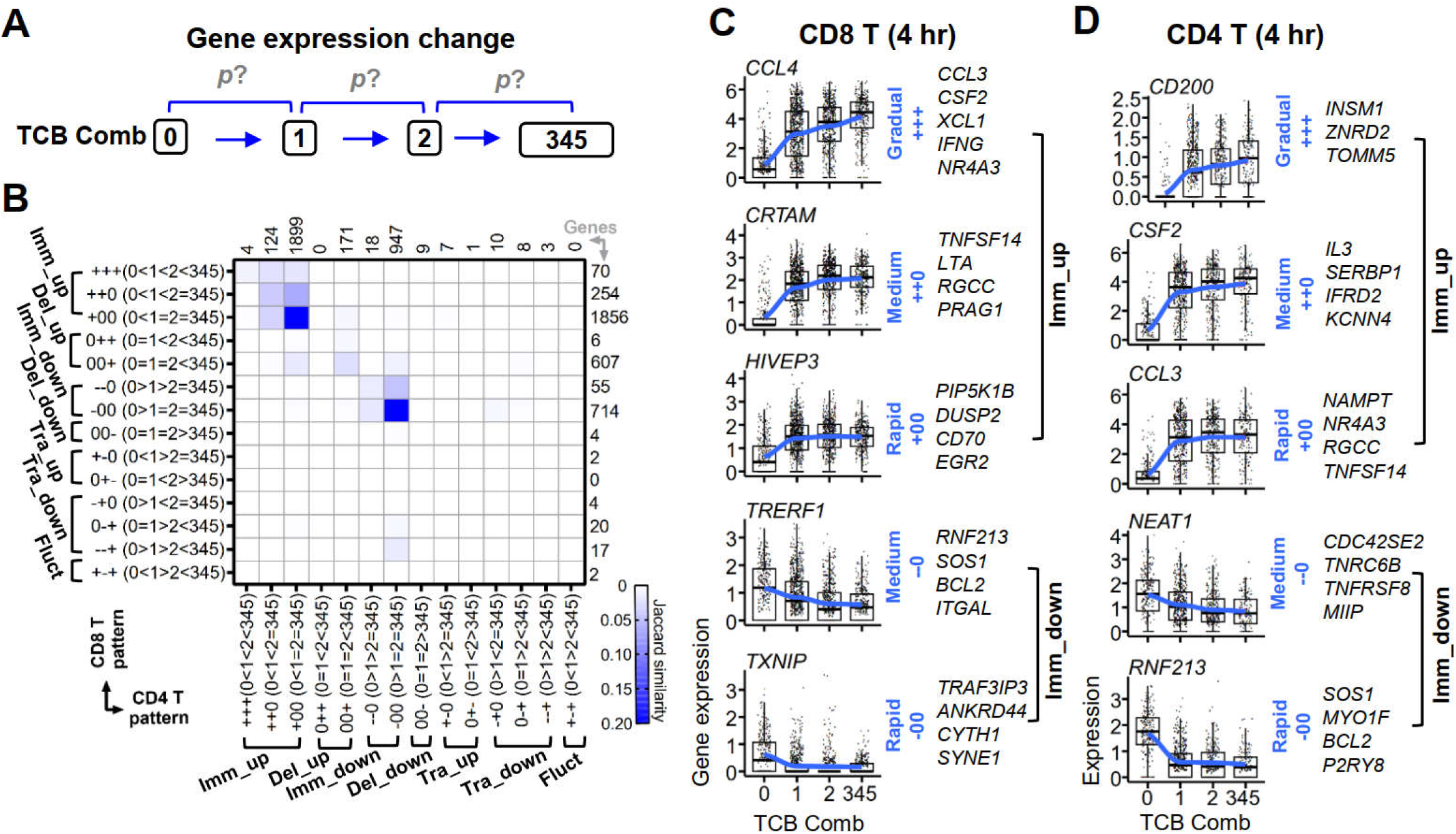
Dynamic gene expression underlies CAR-T serial killing capacity. **(A)** Schematic showing the identification of dynamic gene expression patterns that were associated with the acquisition of serial killing capacity. Differential gene expression between two adjacent groups was tested using Wilcoxon method with Holm correction. **(B)** Gene expression patterns and gene numbers that passed significance test (Padj <0.05). The symbol “+”, “−”, and “0” indicated upregulated, downregulated, and unaltered gene expression between two groups, correspondingly. “Imm”, “Del”, “Tra”, and “Fluet” were short for “Immediate”, “Delayed”, “Transient”, and “Fluctuating” respectively. **(C-D)** Representative plots showing the expression dynamics of selected genes that were immediately up- or downregulated with increasing sequential TCB capture. Gene expression trends were fitted with loess regression, and classified into “Rapid”, “Medium”, and “Gradual” based on the kinetics to reach their highest or lowest expression level.

## DISCUSSION

A multitude of strategies have been deployed to improve the cytotoxic cells’ effectiveness for better disease treatment or prophylaxis. Nevertheless, there remains a significant gap in our knowledge about their functional heterogeneity, and molecular underpinnings that may be manipulated to improve therapeutic outcomes among broader patient groups of diverse genetic background. In this study, we have described the development of DynaKiller-Scan, a robust, high-throughput single-cell multimodal analysis platform designed to quantify the dynamic killing behavior of cytotoxic lymphocytes and link it to molecular programs through integrated transcriptomic profiling.

Fundamentally differing from time-lapse imaging-based identification, single-cell retrieval, and transcriptional profiling of serial killers^9^, DynaKiller-Scan leverages trogocytosis-mediated transfer of differential DNA barcodes from target cells to killer cells, and distinguishes killing dynamics via sequencing-based readout of combinatory target barcode capture. Thus, DynaKiller-Scan requires only simple co-culture of pooled effector-target cells, enabling high scalability and compatibility. In our study, DynaKiller-Scan has captured previous findings based on live-cell imaging^35^ While CD4^+^ CAR-T cells showed slower initial target cell killing relative to CD8^+^ CAR-T cells, their cytotoxic activity ultimately reached comparable levels over time. Meanwhile, within both CD4^+^ and CD8^+^ CAR-T populations, distinct subpopulations exhibited variable killing dynamics and efficiencies. Moreover, through integrated transcriptional and phenotypic co-profiling, DynaKiller-Scan was able to further uncover important gene regulatory networks driving the kinetic variation among refined CAR-T subsets and identify useful phenotypic markers for selecting superior effector cells.

There is growing evidence showing that both CD4^+^ and CD8^+^ CAR-T cells are potential for tumor therapy^33,34,44^, and their functional persistence *in vivo* is a central topic of durable disease remission. T cell of less differentiation before adoptive transfer was thought to predict better therapeutic outcome^44,45^. These cells, however, may develop into more heterogeneous cell states after infusion in complex disease milieu. A small number of long-persisting CD4^+^ CAR-T cells, as characterized by *GZMA*^high^ *GZMK*^high^ and co-expression of both activation and inhibitory markers, were found in two patients with decade-long leukemia remission after receiving CD19-redirected CAR-T treatment^46^. In our study, we found that while *GZMA*^high^ CD8^+^ CAR-T cells were functionally inferior as an exhaustion state, those *GZMA*^high^ CD4^+^ CAR-T cells with the co-expression of T-cell negative regulator *NLRC3* were extremely function-inhibited. On the other facet, new findings showed that elevated type-II cytokine response (e.g., *IL4*, *IL13*, *IL5*, and *GATA3*) but not conventional type-I cytokine functionality (e.g., *IFNG TNF*, *CSF2*, and *TBX21*) endowed CAR-T cells (regardless of CD4^+^ or CD8^+^ lineage) with functional maintenance *in vivo*, and was correlated with durable leukemia remission^47^. However, evidence about the detailed target-eradicating potential and heterogeneity of these cells remains unavailable for these studies. In this work, we demonstrated that while type-I cytokine secretion was involved in the rapid killing activity of both CD8^+^ and CD4^+^ CAR-T cells, elevated type-II functionality favored the killing maintenance of CD8^+^ CAR-T cells. Meanwhile, for CD4^+^ CAR-T, serial killing was most prominent in cells with a mixed type-I/II cytokine secretion response. In future, when combined with clonality analysis or lineage tracing, it could be feasible to exploit DynaKiller-Scan to simultaneously uncover the functional heterogeneity and cell-fate reprograming of therapeutic T or NK cells in combating refractory diseases.

Single-cell CRISPR (scCRISPR) screening approaches that combine pooled genetic perturbations with single-cell multiomics analysis are powerful in dissecting genotype-phenotype relationships for better therapeutics development^48–50^. Recently, *in vivo* scCRISPR screening in mouse solid tumor model, with a focus on 180 transcription factors, has been described to discover gene circuits underlying CTL differentiation for improved therapeutic efficacy^51^. Nevertheless, like many other studies aimed to find candidate genes to regulate effector cell function^52–54^, it has not been able to incorporate the killing function of individual effector cells during the screening procedures. DynaKiller-Scan, given its unique merit in providing single-cell-resolved killing readout, would offer a scalable platform to empower the current genetic and/or chemical screen methods, particularly, to identify gene circuits contributing to the reinvigoration of T-cell exhaustion/dysfunction state^51,55^.

An additional strength of DynaKiller-Scan is its ability to rapidly map function effector-target pairings in complex cellular environments where multiple effector cell types (e.g., CTLs, NK cells, and bystanders) and diverse targets (e.g., different tumor cell types) coexist. This capability enables systematic dissection of multicellular interactions during immune surveillance and reveals functional specificities that are difficult to capture with existing methods. TCR and their cognate tumor neoantigens form the molecular foundation of TIL- and TCR-T–based immunotherapies, as well as personalized cancer vaccines^4,56^. However, parallel discovery of rare TCR–epitope pairs remains challenging under “library-on-library” screening strategies, which combine thousands of candidate antigens with millions of unrelated TCR clonotypes in a single sample^56,57^. Current approaches to enrich antigen-reactive T cells are time-consuming, labor-intensive, and often fail to resolve true effector–target relationships at single-cell resolution^25,57–60^. By directly linking single-cell cytotoxicity with effector–target pairing in a high-throughput and cost-effective manner, DynaKiller-Scan enables the discovery of physiologically relevant TCR–cognate antigen interactions and supports the development of next-generation immunotherapies.

Nevertheless, DynaKiller-Scan has limitations. First, the inherent instability of DNA barcodes limits the time-course of killing assay, which can be potentially overcome by using well-optimized oligonucleotide (e.g., degradation-resistant aptamer)^61,62^. Second, in scenarios wherein multiple target-cell types are involved, it would be necessary to optimize barcode loading efficiency and minimize potential steric interference with effector-target physiological interactions. Finally, while the current study focused primarily on CAR-T cells, expanding this approach to broader lymphocyte populations (e.g., TCR-T, NK, and γδ T cells) will be essential for generalizing its applicability.

Altogether, the DynaKiller-Scan proves a high-throughput, high-resolution platform for simultaneous functional and molecular profiling of individual cytotoxic lymphocytes. We envision that DyanKiller-Scan will be an important tool for the community of diverse research areas to explore the killing behaviors of cytotoxic cells in different diseases, advancing our knowledge of immune cell function and accelerating the development of new vaccines and therapeutics.

## METHODS

### Primary cells and cell lines

Human anti-CD19 CAR-T cells were generated as previously described^63^. Briefly, apheresis residual blood-derived peripheral blood mononuclear cells (PBMC) were activated by anti-CD3/CD28 Dynabeads (Gibco, 11161D) at a 2:1 bead-to-cell ratio one day before lentivirus transduction to co-express 4-1BB-CD3-zeta anti-CD19 CAR and GFP (CAR.19-4-1BBz-IRES-GFP). The viruses were removed after 3 days post-infection, and expanded for 9-10 more days in RPMI 1640 medium (Gibco, A1049101) supplemented with 10% fetal bovine serum (FBS) (Gibco, 10270106) and 5 ng ml^−1^ IL-2 (Gibco, PHC0021) at an interval of 3-4 days. Human primary NK cells were enriched from PBMC using EasySep human NK cell enrichment kit (STEMCELL Technologies, 19055) and expanded for 9-10 days using the ImmunoCult NK cell expansion kit (STEMCELL Technologies, 100-0711)^22^. Both CAR-T and pre-expanded NK cells were aliquoted and stored in liquid nitrogen. Human B-cell precursor leukemia cell line NALM6 (ATCC, CRL-3273) and chronic myeloid leukemia cell line K562 (ATCC, CCL-243) were maintained in RPMI 1640 medium with 10% FBS.

### Cell-surface modification with fluorochromes

NALM6 or K562 cells were stained with Sulfo-NHS-LC-Biotin (NHS-biotin; Thermo Scientific, A39257) from 0-500 µM as indicated at 8 million (M) cells per milliliter (ml) in PBS (pH 7.4). The cells were incubated at room temperature (RT) for 30 min before extensively washed with phosphate buffered saline (PBS) containing 2% FBS to remove free biotin. The cells were stained with fluorochrome-labelled streptavidin (Strep-Fluo) as indicated at 4°C for 30 min: 2 µl Strep-PE (BioLegend, 405203; stock 0.2 µg µl^−1^), or 2 µl Strep-APC (BioLegend, 405207; stock 0.2 µg µl^−1^), 1 µl Strep-BV421 (BioLegend, 405225; stock 0.5 µg µl^−1^), or 4 µl Strep-BV785 (BioLegend, 405249; stock 0.1 µg/µl), correspondingly. The staining was further blocked with D-Biotin at RT for additional 15 min, and washed three times with 2%FBS.

### DynaKiller-Scan with multiparametric flow cytometry

CAR-transduced T cells (∼20% GFP^+^ CAR-T) were recovered overnight and activated with anti-CD3/CD28 DynaBeads (1:1 bead:cell ratio) in the presence of 5 ng ml^−1^ IL-2 for additional 3-4 day. For single-color DynaKiller-Scan, NALM6 cells (“N6”) were labelled with Strep-PE and co-cultured with effector T cells at a 1:4:1 CAR-T:Non-CAR-T:N6 ratio. To distinguish CAR-T and NK mediated specific trogocytosis, the co-culture system composited of 1:1:45 CAR-T:NK:N6 or CAR-T:NK:K562. For multi-color DynaKiller-Scan, NALM6 cells (“N6”) were differentially labelled with Strep-PE, Strep-BV421, Strep-BV786 as described above before equally pooled and co-cultured with T cells in 12-well cell culture plate in 2 ml culture medium as follows: total T: N6 = 0.15 M : 0.05 M, 0.05 M : 0.15 M, or 0.05 M : 0.45 M. Where indicated, anti-CD107a-AF647 (BioLegend, 328612; stock 0.4 µg µl^−1^) were included at final concentration of 1 µg ml^−1^. After 4 hr, cells were incubated at 4oC for 10 min before collection.

The cells were spun down, washed, and analyzed by flow cytometry. To calculate the number of target cells remained after co-culture, precision count beads (Biolegend, 424902) were spiked in the cell suspension before flow cytometry analysis.

### Cell-surface modification with DNA oligonucleotides

NALM6 cells were modified with 4, 20, and 100 µM NHS-biotin at RT for 30 min before conjugated with 2 µl Strep-PE (stock 0.2 µg µl^−1^) for additional 30 min at 4°C. The cells were washed twice with 2% FBS and further stained with 400 pmol biotin-oligo conjugate at RT for 30 min. In some experiments, the biotinylated cells were immediately stained with 1 µl TotalSeq™-A0951 Strep-PE (BioLegend, 405251; stock 0.5 µg µl^−1^) at 4°C for 30 min. To confirm the successful labelling of biotin-oligo, the cells were stained with 200 pmol FAM-polyT probe (stock 200 µM) or 100 pmol Cy5-polyT probe (stock 200 µM) in 2% FBS at RT for 30 min, and detected by flow cytometry.

### Quantifying the trogocytosis of DNA label via quantitative PCR (q-PCR)

NALM6 cells were modified with 20 µM NHS-biotin at RT for 30 min before conjugated with Strep-PE for additional 30 min at 4°C. The cells were stained with 400 pmol biotin-oligo at RT for 30 min and blocked with D-Biotin (stock 0.25 ug/ul) for 15 min. The cells were washed twice with 2% FBS before co-cultured with T cells (∼ 20% CAR-T cells) in 12-well cell culture plate in 2 ml culture medium as follows: total T: N6 = 0.15 M : 0.05 M. Anti-CD107a-AF647 (BioLegend, 328612; stock 0.4 µg µl-1) were included at final concentration of 1 µg ml^−1^. After 4 hr, cells were incubated at 4°C for 10 min before collection. T cells were washed twice with 2%FBS and sorted into CD107a^high^ PE^high^ group (*16 single cells) and those remained counterpart (*16 single cells) (gating based on a total of ∼ 0.5% background capture by Non-CAR-T cells). Single-cell q-PCR was performed to test the sensitivity and specificity of oligo trogocytosis following method described previously^64^.

### DynaKiller-Scan with multimodal sequencing

Strep-PE^+^ NALM6 cells were split into six groups and barcoded with biotin-oligo, correspondingly. The differentially barcoded NALM6 were co-cultured with T cells (CAR-T : Non-CAR-T : total N6 = 0.01 M : 0.01 M : 0.45 M) in 12-well cell cultured plate (Costar, 3513) in 2 ml culture medium. CD107a-AF647 were included at a final concentration of 1 µg ml^−1^. At the time point of 1 hr and 4 hr, cells were collected, and excessive NALM6 targets were removed using EasySep™ Human CD19 Positive Selection Kit II (STEMCELL Technologies, 17854). The recovered total T cells were resuspended in 2% FBS. For each time point, according to the background capture of Non-CAR-T cells, CAR-T counterpart was sorted into PE^high^ CD107a^high^ and those remained population. The sorted CD107a^high^ PE^high^ CAR-T (1 hr), CD107a^low^ PE^low^ CAR-T (1 hr), CD107a^high^ PE^high^ CAR-T (4 hr), and CD107a^low^ PE^low^ CAR-T (4 hr) were individually barcoded with 0.5 µg β2M/CD298-targeting TotalSeq-A hashtag 1 (BioLegend, 394601), hashtag 2 (BioLegend, 394603), hashtag 3 (BioLegend, 394605), and hashtag 4 (BioLegend, 394607) respectively, together with Fc-blocking agents (BioLegend, 422302) at 4°C for 30 min. Cells were extensively washed with 2% FBS before subjected to cell-bead emulsion generation using 10x Chromium Controller (10x Genomics).

### Sequencing library preparation

Single-cell barcoded DNA libraries, including complementary DNA (cDNA) derived from mRNA and protein libraries representative of trogocytosis-derived target-cell barcodes, cell surface protein markers and sample-indication hashtags, were generated using 10x Genomics single-cell 3’ reagent kit (v3.1 Chemistry) as described previously^64^. Briefly, following reverse transcription and sample clean-up, 35 µl of the eluted DNA sample was pre-amplified as follows: 98°C 3 min, 12 cycles of: 98°C 15 s, 63°C 20 s, and 72°C 1 min; Then an extension step of 1 min at 72°C. Protein libraries and cDNA library were separated following a double size selection step with SPRIselect reagent (Beckman Coulter). The amplified cDNA product was subjected to final gene expression library construction according to the instructions with the kit. Part of the protein library product (5 µl) was used for the preparation of sequencing libraries corresponding to surface marker and captured target-cell barcodes while additional part (5 µl) was used for the construction of sequencing library of sample hashtags using KAPA HiFi Master Mix (Roche). The size range and concentration of final library constructions was verified by Agilent 2100 Bioanalyzer.

### Sequencing data pre-processing

The libraries were pooled and sequenced in Illumina NovaSeq X Plus platform. Raw FASTQ data was processed to call cell barcodes and unique molecular identifier (UMI) of cDNA library using Cell Ranger v5.0.1 (10x Genomics) with default parameters. The filtered cell barcodes were used to find individual cell barcode-matched DNA tags including sample hashtags, surface proteins-targeted antibody-derived tags (ADT), and captured target-cell barcodes (TCB) using CITE-seq-Count (v1.4.5; doi: 10.5281/zenodo.2590196) with a Hamming distance of 1 for both cell barcodes calling and UMI counting. The raw count matrices were imported into Seurat package (v5.0.2)^65^. Dead cells were removed by filtering out cells with >12% mitochondrial transcript counts. Cells of high-quality sequencing output, identified by gene features >200 but <10,000, and total gene counts <100,000 were kept for further data quality control. The read counts of mRNA-derived cDNA were log-transformed with LogNormalize method, while sample hashtags were normalized using a centered log ratio transformation (CLR). Cells were demultiplexed based on relative hashtag abundance by using HTODemux() function with default parameters. To further remove cell doublets or any unwanted contamination potentially derived from unknown libraries during pooled sequencing, only cells with apparent hashtag staining (hashtag 1 >1.5, hashtag 2 >3.0, hashtag 3 >2.0, and hashtag 4 >3.0 respectively) were included for downstream analysis. Cell cycle phase scores were calculated with CellCycleScoring() function using a list of canonical mitosis-associated genes, and regressed during principal component analysis (PCA).

### Processing of ADT read counts

One million of dropped cell barcoded (containing empty droplets) were randomly recovered from the Cell Ranger unfiltered output to evaluate the ambient level of individual ADT (cell-unbound ADT). Qualified empty droplets, compared to cell-containing droplets, were identified by cDNA size <2, and protein size >2 but <3. Thus, for each ADT in each individual cell, the cell-bound ADT were calculated by subtracting total read counts of that ADT from its average reads in empty droplets. In addition, to compensate antibody isotype-introduced staining variation at cell-level, the cell-bound ADT were further subtracted from its isotype staining within the same cell. These processed ADT read counts were normalized using a centered log ratio transformation (CLR) and imported into Seurat object for downstream multimodal analysis.

### Processing of TCB read counts

As shown in Supplementary Fig. 5, there was substantial transcriptomic difference towards cell activation, polyfunctional secretion, cytotoxicity of the sorted CD107a^hi^ PE^hi^ CAR-T cells over CD107^low^ PE^low^ counterpart, suggesting that most killers could be found within the CD107a^hi^ PE^hi^ group while non-killers would be within the CD107a^low^ PE^low^ group. Thus, unsupervised cell clustering based on global gene transcription was performed for each time point to find the main cell cluster (cluster 4 for 1 hr and cluster 2 for 4 hr) highly enriched in the CD107a^low^ PE^low^ group (Supplementary Fig. 6a). This cell cluster was used for the initial threshold setting of each TCB background capture (<99% quantile), correspondingly (Supplementary Fig. 6b). The read counts of TCB were centered log-ratio transformed. By this, we were able to roughly categorize the combinatorial trogocytosis of TCB from “Comb 0” to “Comb 6”.

Next, we were wondering if there could be some killer cells falsely identified as negative TCB capture due to detection sensitivity. To verify this possibility, unsupervised transcriptome clustering was re-performed for those cells of negative TCB capture (previously identified as “Comb 0”) for each individual time-point. As anticipated, many of the cells from the sorted CD107a^hi^ PE^hi^ CAR-T were transcriptionally distinguishable from from the CD107^low^ PE^low^ CAR-T (Supplementary Fig. 7a,b). Therefore, these cells were re-annotated as cells of undetermined TCB capture (“UD”) and true negative capture (“Comb 0”).

### T-cell lineage and subset annotation

Via combined analysis of the relative expression of lineage-specific genes (*CD4*, *CD8A*, and *CD8B*) and surface protein markers (CD4 and CD8a), T cells from individual time point were categorized into CD4^+^ T, CD8^+^ T, double positive (DP) T, and double negative (DN) T cells (Supplementary Fig. 9). DoubletFinder (v2.0.6)^66^ was used to check the potential doublet contamination among the identified T-cell lineages. To characterize subtle T populations, we utilized the reciprocal PCA (RPCA) method (implemented in the Seurat package) to integrate CD4^+^ T or CD8^+^ T cells from different time points, correspondingly, based on their global gene expression profiles. Cell cycle effect was calculated with CellCycleScoring() function, and regressed during data scaling and PCA performing. Differential gene expression analysis by Wilcoxon rank-sum test using FindAllMarkers() function, setting min.pct =0.20 and logfc.threshold =0.5, was performed to find CD8^+^ or CD4^+^ cluster-specific marker genes. Genes passing Bonferroni-adjusted significance test (*P*adj <0.05) were selected for downstream analysis. Jaccard similarity was performed to find the most likely matched cell subsets across CD4^+^ T and CD8^+^ T cells.

### Gene set enrichment analysis (GSEA)

Differential gene expression analysis by Wilcoxon Rank Sum test was performed using FindAllMarkers() with the settings of min.pct =0.20 and logfc.threshold =0. Whole gene lists generated were used for GSEA^67^ which has been embedded in clusterProfiler (v4.10.1)^68^. Where indicated, the curated hallmark gene sets or GO biological processes were downloaded from the Molecular Signatures Database (MSigDB, v2025.1) for gene set prediction^67^. Enriched gene sets were ranked by normalized enrichment scores (NES) and adjusted *P* value (*P*adj <0.05).

## Data availability

Curated hallmark gene sets and GO biological processes are publicly available from Molecular Signatures Database (MSigDB, v2025.1). The list of genes encoding transcription factors (TFs) were from Lambert et al (v1.01; https://humantfs.ccbr.utoronto.ca/index.php)^69^. The gene lists for secreted proteins and those non-secreted or non-TFs were retrieved from Human Protein Atlas (v24.0; https://www.proteinatlas.org/humanproteome/tissue/secretome)^70^. Single-cell sequencing data and code to reproduce the analyses in this work are available on request.

## Acknowledgements

This work was supported by the National Research Foundation (NRF-000510-00-00) and Ministry of Education of Singapore (MOE-T2EP30224-0020). We thank Professor Michael Birnbaum (MIT) for providing the anti-leukemia CAR-T cells used in this study.

## Author Contributions

T.W and L.F.C conceived and designed the study. T.W performed the experiments and data analysis with input from L.F.C. T.W and L.F.C wrote the manuscript. All authors approved the submission.

