## Supplementary information for "DynaKiller-Scan: Multimodal characterization of single-cell killing dynamics via combinatorial encoding of target lysis"

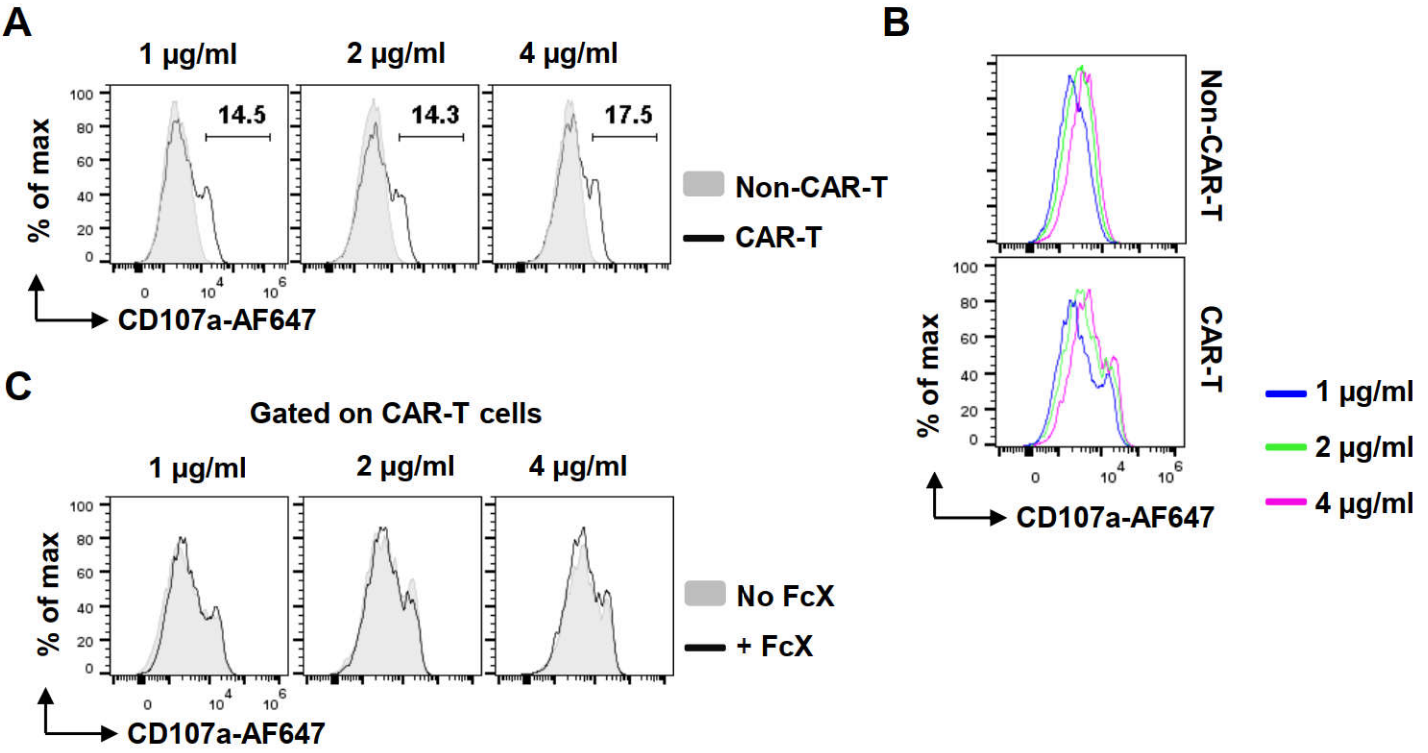


**Supplementary Figure 1. Optimization of anti-CD107a concentration used in co-culture assay**

(**A**) Different concentrations of anti-CD107a-AF647 was included during co-culture assay. After 4 hr, the staining intensity of CD107a on CAR-T versus Non-CAR-T cells were detected by flow cytometry. (**B**) Shown were histograms of CD107a staining gated on CAR-T and Non-CAR-T cells respectively. (**C**) Effect of Fc receptor pre-blocking on the detection of anti-CD107a-AF647 during co-culture.

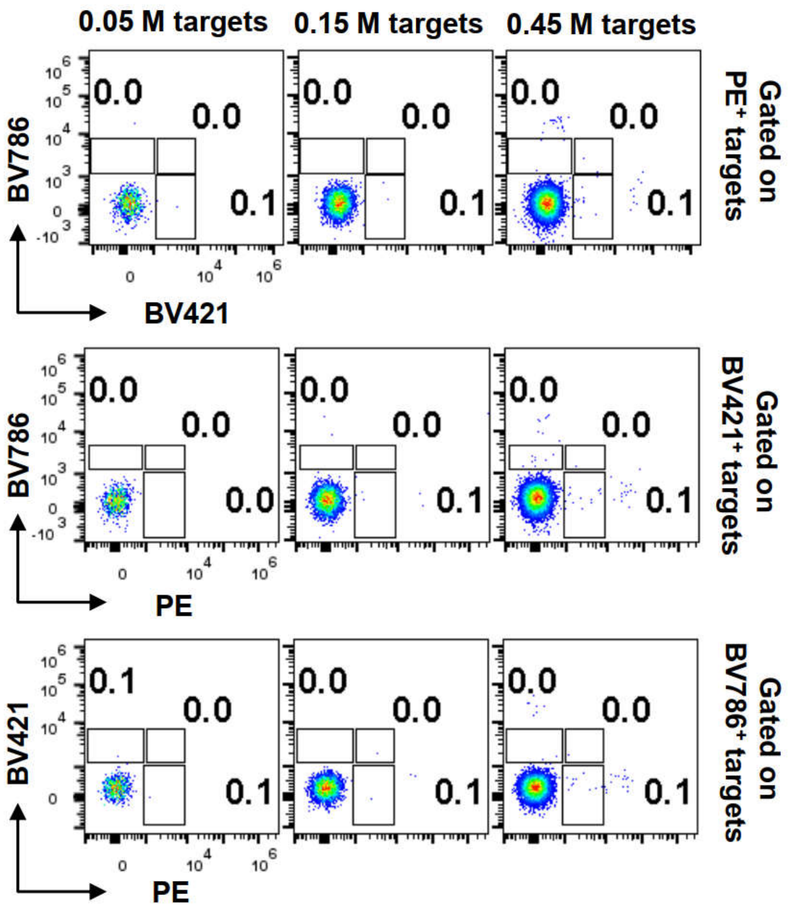


**Supplementary Figure 2. Cross-capture of labelled dyes among NALM6 cells**

Differentially colored NALM6 cells were seeded onto 12-well cell culture plate at indicated density, and co-cultured with CAR-T cells. After 4 hr, cross capture of labelled dyes between NALM6 cells was detected by flow cytometry.

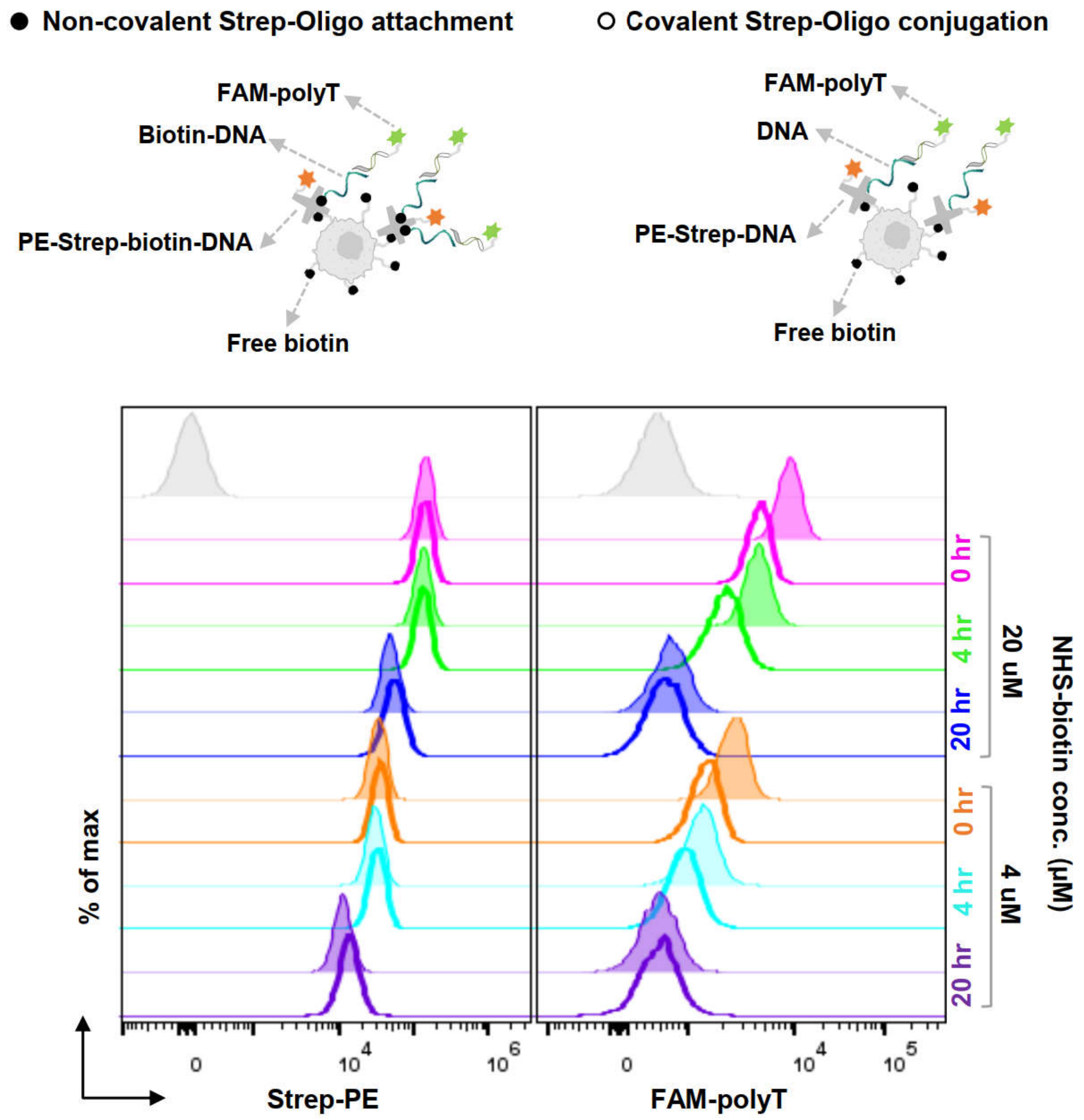


### Supplementary Figure 3. Comparison of DNA barcode loading approaches

Biotinylated NALM6 cells were stained with Strep-PE before being loaded with biotin-conjugated DNA barcodes (non-covalent attachment, solid symbol), or directly modified with Strep-PE-DNA complex (covalent attachment, open symbol). The stability of on-cell Strep-PE and DNA barcode (measured by FAM-polyT probe binding) in culture medium was assessed over time.

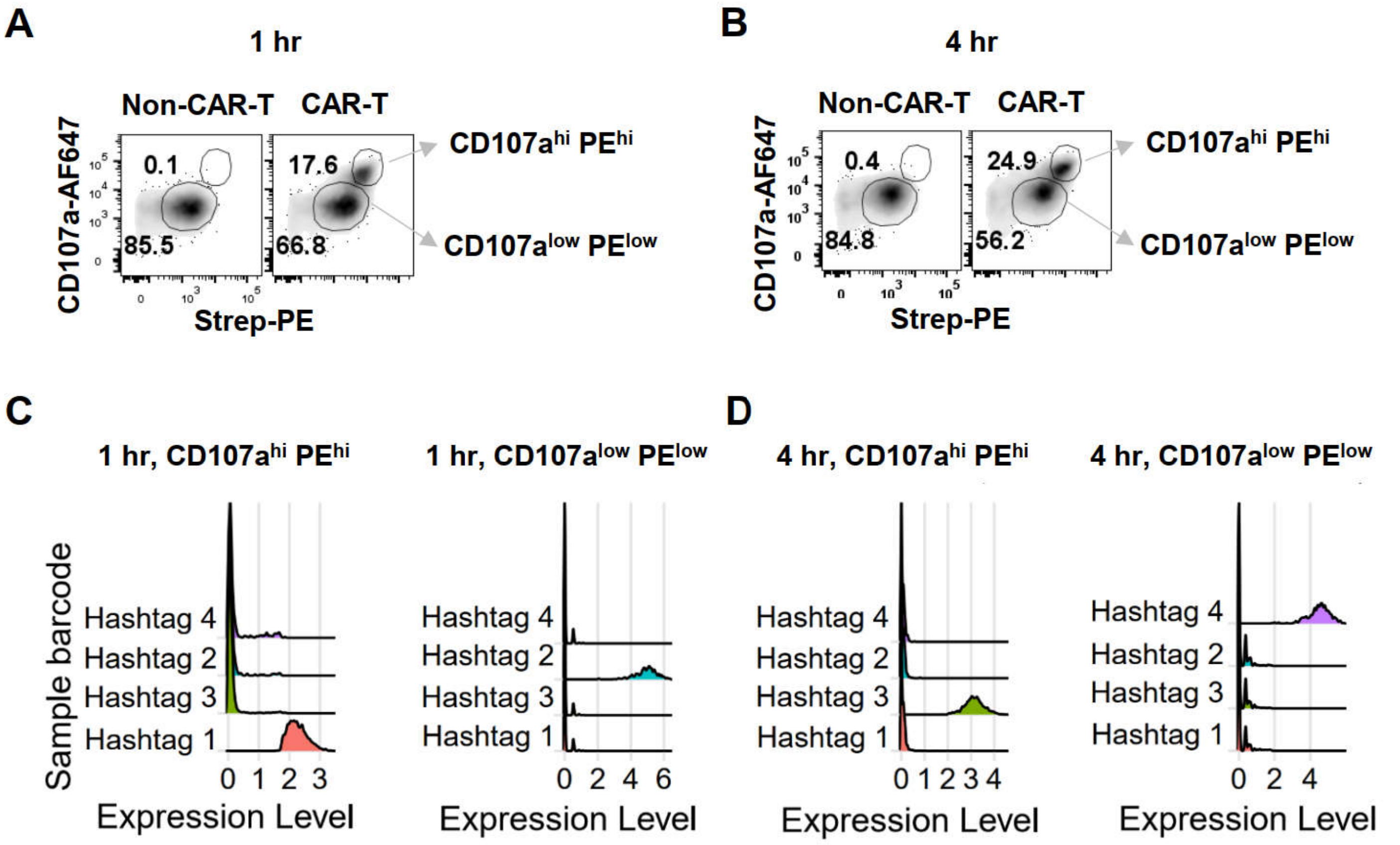


**Supplementary Figure 4. Cell sorting, sequencing, and demultiplexing**

(**A**-**B**) After 1 or 4 hr co-culture, excess NALM6 cells were depleted using immunomagnetic selection, and remaining CAR-T cells were sorted into CD107a$^{hi}$ PE$^{hi}$ and CD107a$^{low}$ PE$^{low}$ populations. (**C**-**D**) Sample demultiplexing based on normalized hashtag abundance of each sorted cell population.

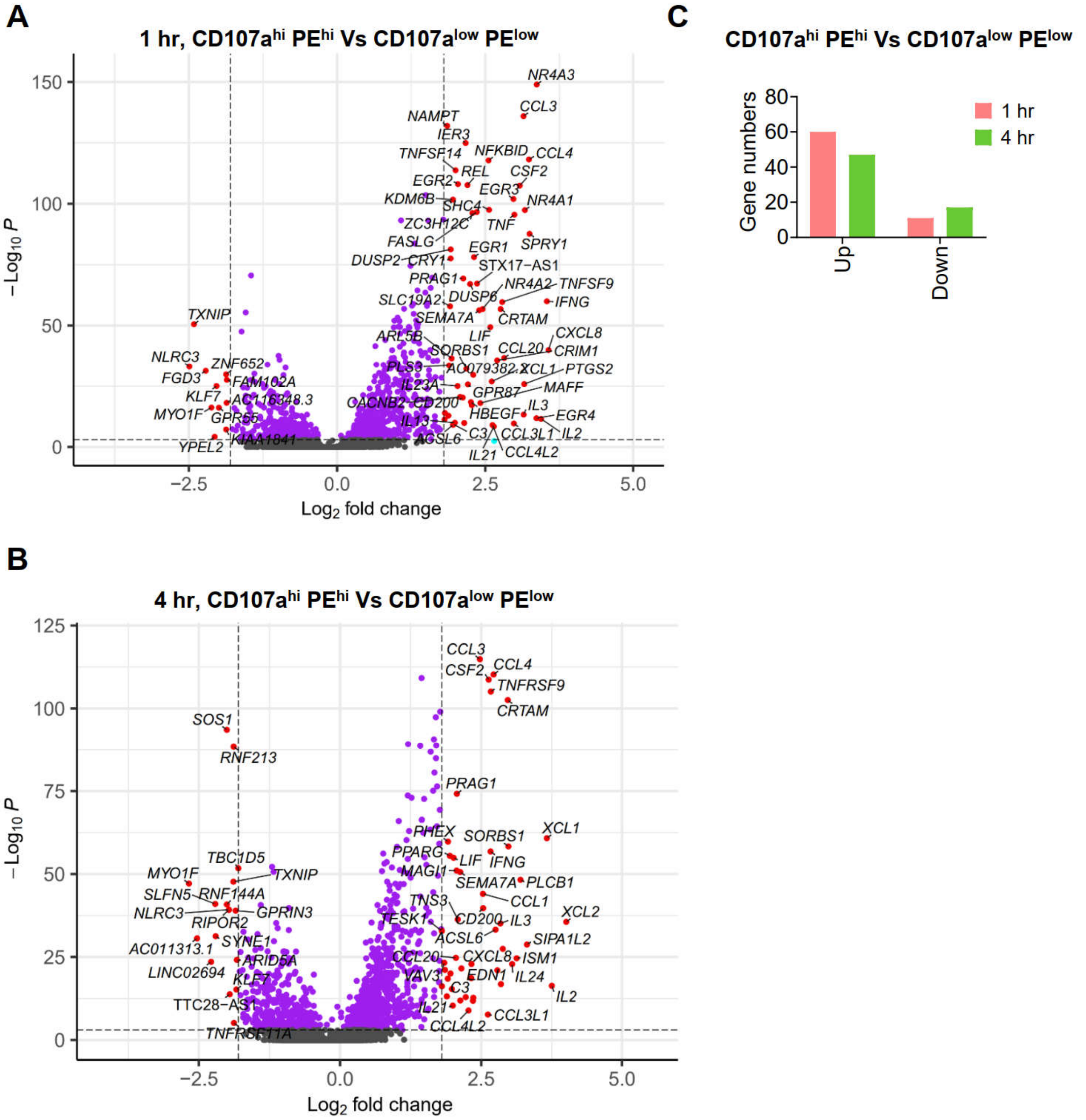


**Supplementary Figure 5. Global gene expression difference of sorted cell populations**

Transcriptional difference between sorted CD107a$^{hi}$ PE$^{hi}$ and CD107a$^{low}$ PE$^{low}$ CAR-T cells. Differentially expressed genes were calculated using FindMarkers() with min.pct =0.2. Representative genes with Bonferroni-adjusted $P$ < 0.05 (two-sided) and log2(fold change) ≥0.5 or ≤−0.5 were highlighted in red in volcano pots (**A**-**B**), and total gene numbers within the threshold were counted (**C**).

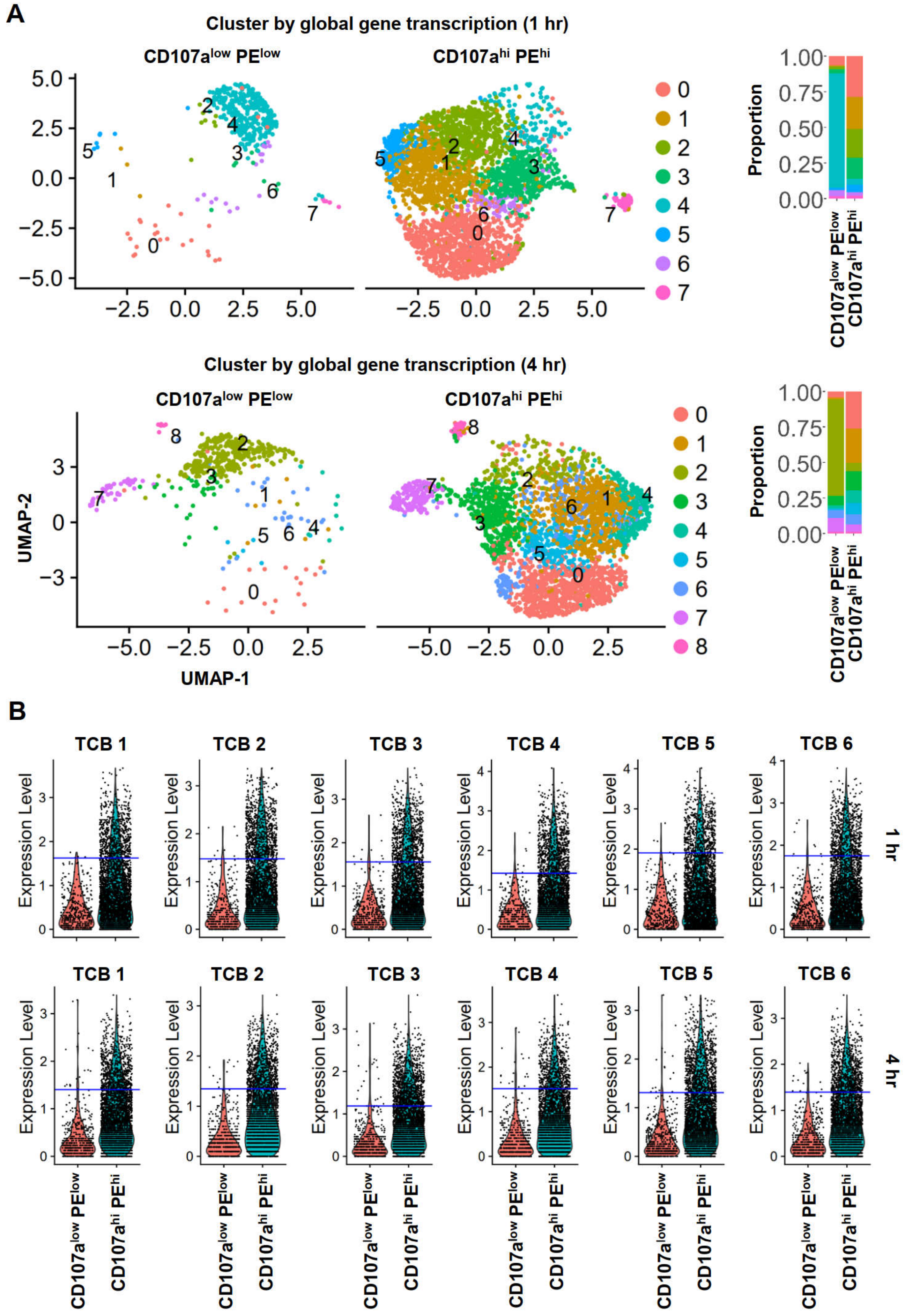
A
Cluster by global gene transcription (1 hr)
CD107a^low PE^low
CD107a^hi PE^hi
Proportion
Cluster by global gene transcription (4 hr)
UMAP-1
UMAP-2
B
TCB 1
TCB 2
TCB 3
TCB 4
TCB 5
TCB 6
Expression Level
1 hr
4 hr

**Supplementary Figure 6. Determination of the background capture of target cell barcodes (TCB)**

(**A**) Unsupervised cell clustering based on global gene expression for samples of individual time points. (**B**) 99% percentile of each TCB captured by the cell cluster 4 (for 1 hr) and 2 (for 4 hr) in **A** was used for the initial threshold setting (blue line) of the TCB background capture.

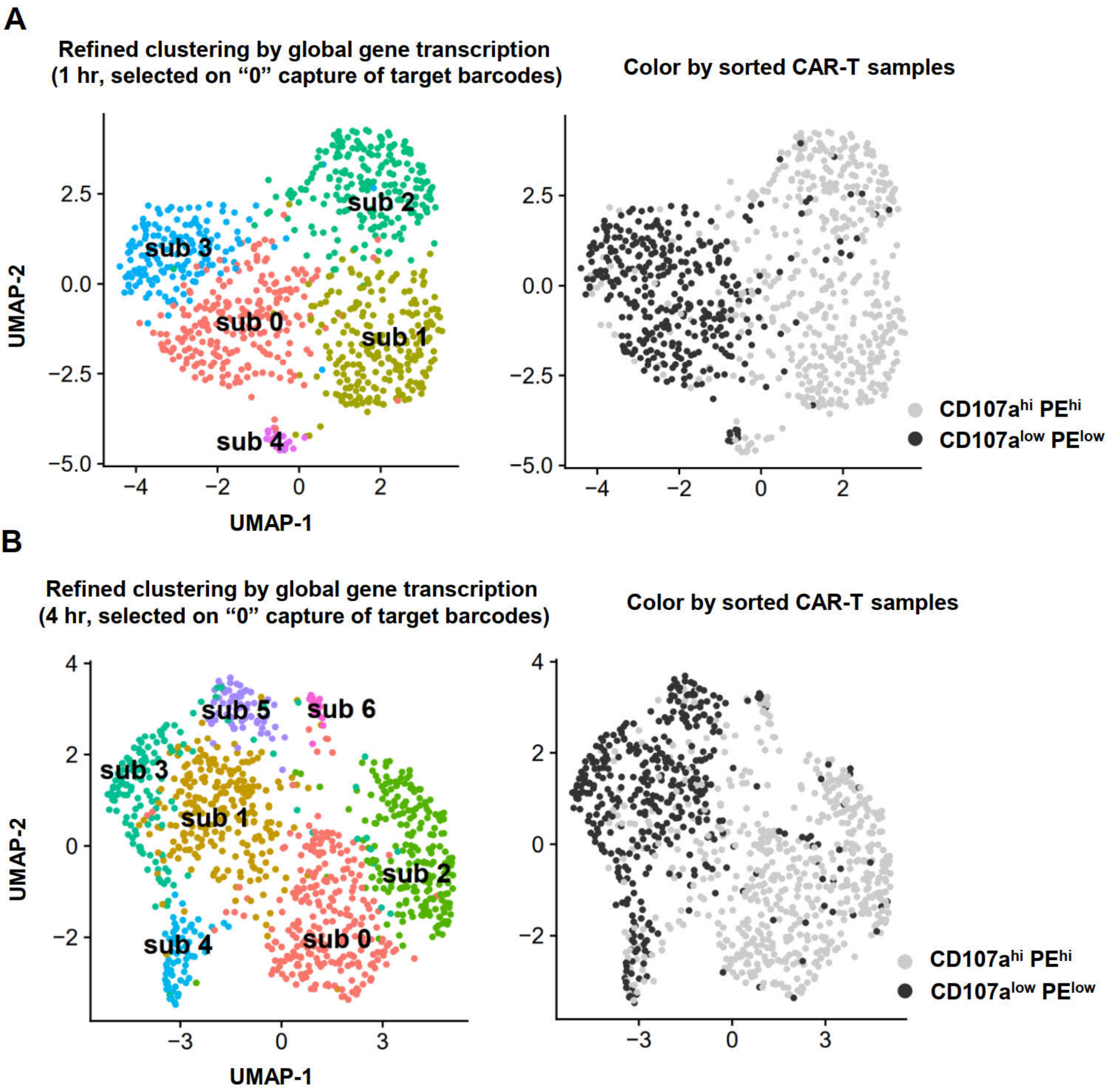


**Supplementary Figure 7. Refined categorization of negative TCB capture**

(**A-B**) Cells initially identified as negative TCB capture (those below cutting threshold in Supplementary Figure 6b) were re-clustered by their transcriptomic variation (left panel), and CD107a$^{hi}$ PE$^{hi}$ and CD107a$^{low}$ PE$^{low}$ CAR-T cells in the re-clustered cell topological structure was shown as well (right panel). Consequently, for 1 hr (**A**), the subcluster "1" and "2" were identified as undetermined ("UD") TCB capture while those remaining being “0” TCB capture. Likewise, for 4 hr (**B**), the subcluster "0" and "2" were identified as undetermined ("UD") TCB capture while those remaining being “0” TCB capture.

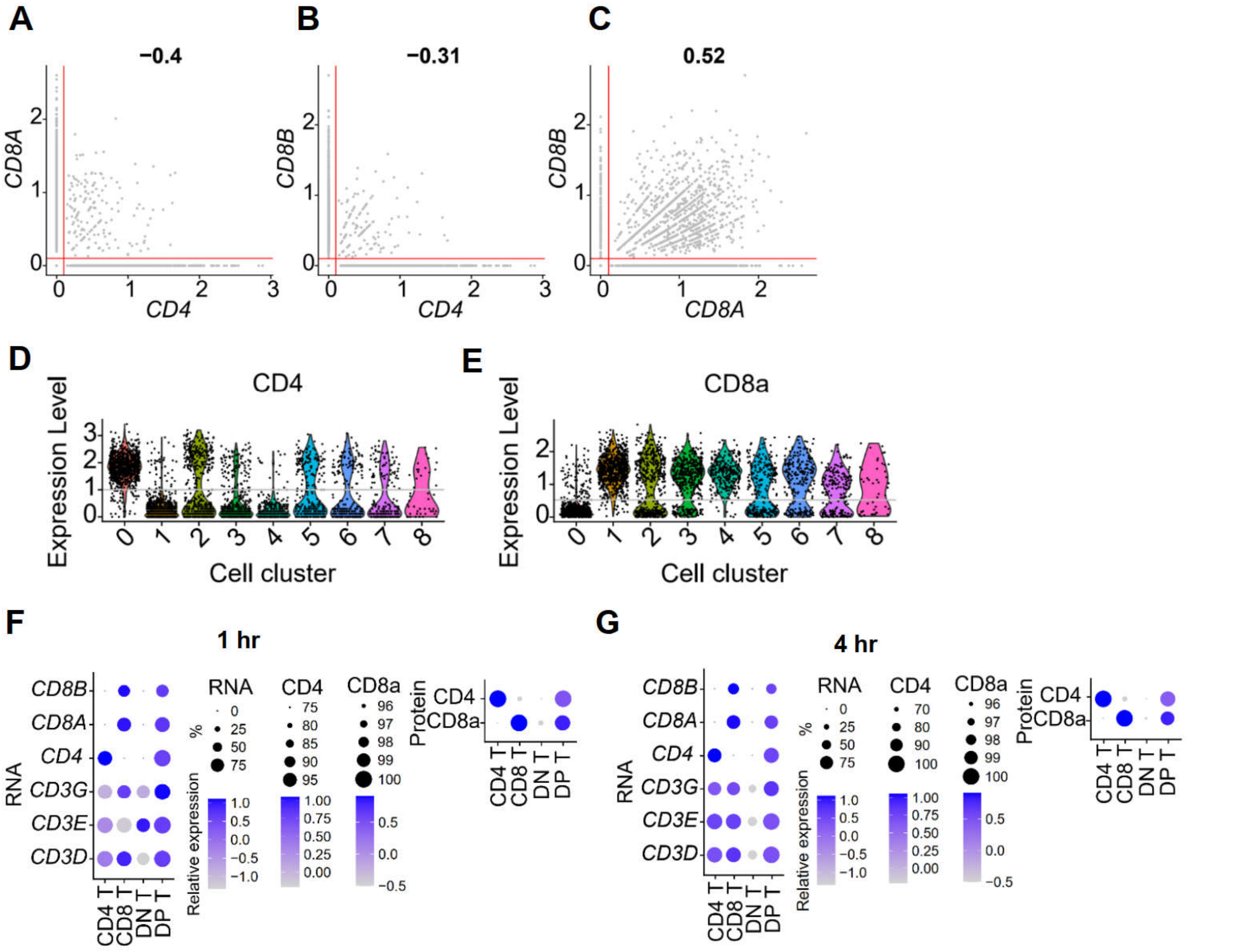


**Supplementary Figure 8. Identification of main T cell lineages**

(**A**-**C**) Representative scatter plots showing the relative expression of T-cell lineage-specific genes including *CD4*, *CD8A*, and *CD8B* (for 4 hr). Threshold (intercept at 0.2) to separate positive and negative cell groups was indicated in red line. (**D**-**E**) Relative expression of T-cell lineage-specific protein markers including CD4 and CD8a among identified cell clusters (for 4 hr). Threshold to separate positive and negative cell groups was indicated by the grey line. (**F**-**G**) The relative expression of lineage-specific genes (*CD3D*, *CD3E*, *CD3G*, *CD4*, *CD8A*, and *CD8B*) and proteins (CD4 and CD8a) in the $CD4^{+}$ T, $CD8^{+}$ T, $CD4^{+}$ $CD8^{+}$ double-positive T (DP T), and $CD4^{-}$ $CD8^{-}$ double-negative T (DN T) populations.

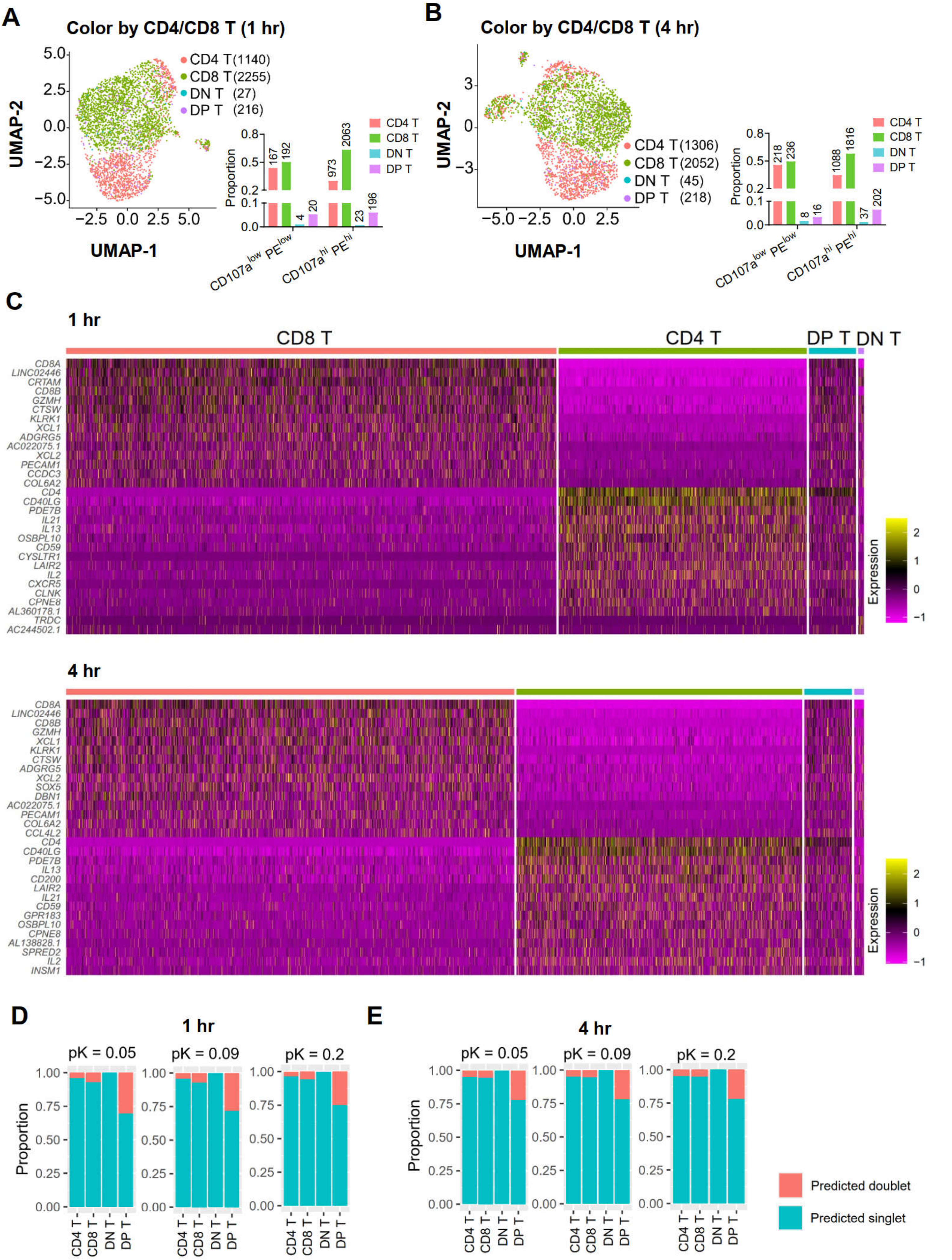

A
Color by CD4/CD8 T (1 hr)
CD4 T (1140)
CD8 T (2255)
DN T (27)
DP T (216)
UMAP-1
UMAP-2
Proportion
CD107a^low PE^low
CD107a^hi PE^hi
B
Color by CD4/CD8 T (4 hr)
CD4 T (1306)
CD8 T (2052)
DN T (45)
DP T (218)
C
1 hr
CD8 T
CD4 T
DP T
DN T
Expression
4 hr
D
1 hr
pK = 0.05
pK = 0.09
pK = 0.2
E
4 hr
Predicted doublet
Predicted singlet

**Supplementary Figure 9. Differences in transcriptional profiles of identified T cell lineages**

(**A**-**B**) Topological distribution of identified T cell lineages in the unsupervised transcriptomics-based cell clustering (UMAP plots), and composition change among sorted CD107a$^{hi}$ PE$^{hi}$ and CD107a$^{low}$ PE$^{low}$ CAR-T cells (histogram plots). (**C**) Heatmaps showing the top differentially expressed genes between CD8$^{+}$ T, CD4$^{+}$ T, DP T, and DN T CAR-T cells. (**D**-**E**) Potential contamination of cell doublets among the identified T-cell lineages was checked with DoubletFinder (v2.0.6).

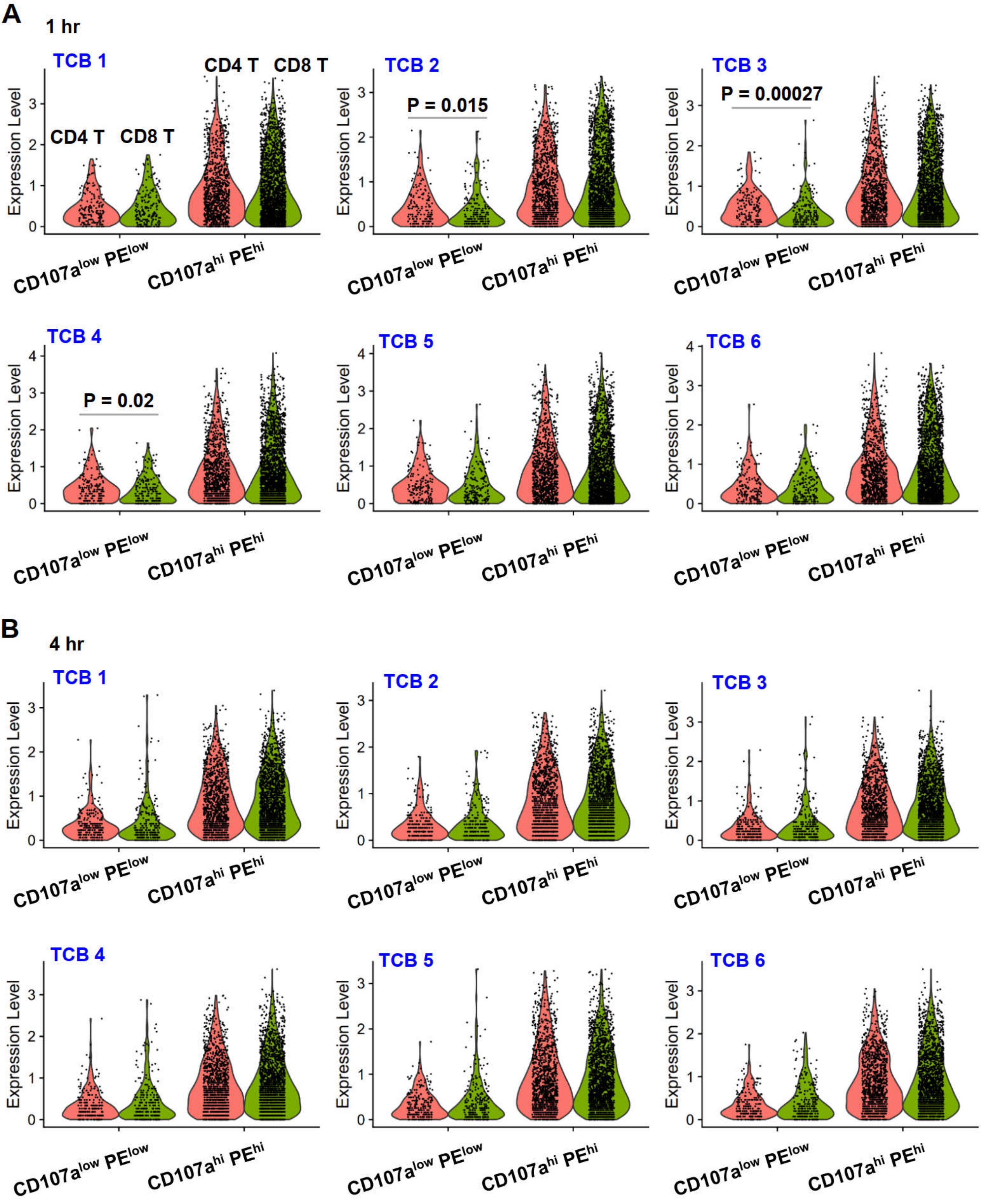


**Supplementary Figure 10. Comparable abundance of each TCB captured by CD4⁺ and CD8⁺ CAR-T cells**

The read counts of TCB were centered log-ratio transformed. Shown were the relative abundance of each TCB in identified $CD4^+$ (red) and $CD8^+$ (green) CAR-T cells among the sorted $CD107a^{hi}$ $PE^{hi}$ or $CD107a^{low}$ $PE^{low}$ CAR-T population. The significance of TCB capture between $CD4^+$ and $CD8^+$ CAR-T cells was tested using Wilcoxon method with Holm correction. Only significant comparisons ($P$ <0.05) were marked.

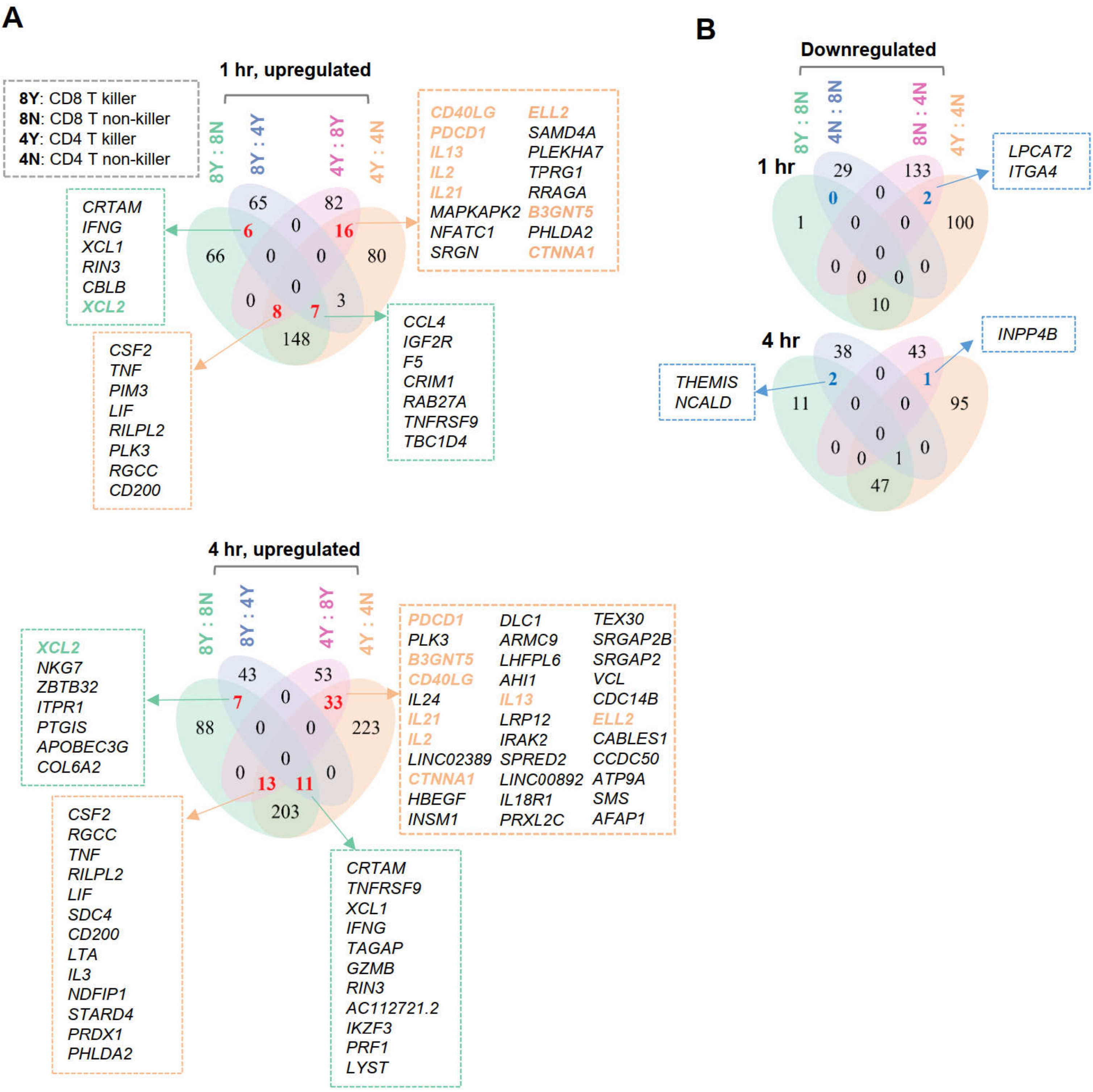


**Supplementary Figure 11. Identification of killing-associated gene expression distinguishing CD8⁺ from CD4⁺ CAR-T killers**

CD8⁺ or CD4⁺ CAR-T killing associated genes that were either upregulated (**A**) or downregulated (**B**) Mutual gene expression difference was calculated using FindMarkers() with the settings of logfc.threshold =0.5 and min.pct =0.20. Genes with Bonferroni-adjusted *P* <0.05 (two-sided) were kept and shown by Venn diagrams.

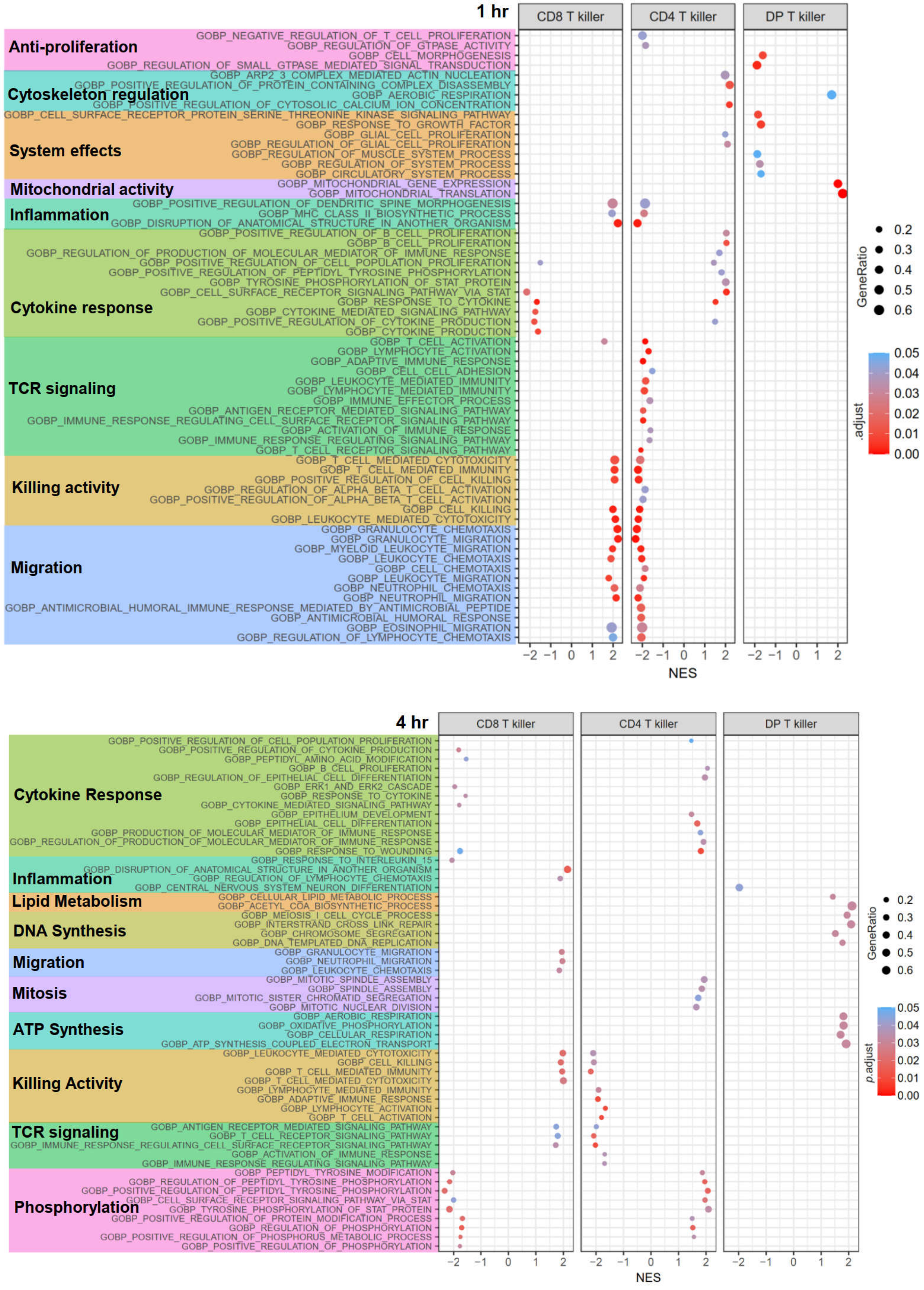
1 hr
CD8 T killer
CD4 T killer
DP T killer
Anti-proliferation
GOBP_NEGATIVE_REGULATION_OF_T_CELL_PROLIFERATION
GOBP_REGULATION_OF_GTPASE_ACTIVITY
GOBP_CELL_MORPHOGENESIS
GOBP_REGULATION_OF_SMALL_GTPASE_MEDIATED_SIGNAL_TRANSDUCTION
Cytoskeleton regulation
GOBP_ARP2_3_COMPLEX_MEDIATED_ACTIN_NUCLEATION
GOBP_POSITIVE_REGULATION_OF_PROTEIN_CONTAINING_COMPLEX_DISASSEMBLY
GOBP_AEROBIC_RESPIRATION
GOBP_POSITIVE_REGULATION_OF_CYTOSOLIC_CALCIUM_ION_CONCENTRATION
System effects
GOBP_CELL_SURFACE_RECEPTOR_PROTEIN_SERINE_THREONINE_KINASE_SIGNALING_PATHWAY
GOBP_RESPONSE_TO_GROWTH_FACTOR
GOBP_GLIAL_CELL_PROLIFERATION
GOBP_REGULATION_OF_GLIAL_CELL_PROLIFERATION
GOBP_REGULATION_OF_MUSCLE_SYSTEM_PROCESS
GOBP_REGULATION_OF_SYSTEM_PROCESS
GOBP_CIRCULATORY_SYSTEM_PROCESS
Mitochondrial activity
GOBP_MITOCHONDRIAL_GENE_EXPRESSION
GOBP_MITOCHONDRIAL_TRANSLATION
Inflammation
GOBP_POSITIVE_REGULATION_OF_DENDRITIC_SPINE_MORPHOGENESIS
GOBP_MHC_CLASS_II_BIOSYNTHETIC_PROCESS
GOBP_DISRUPTION_OF_ANATOMICAL_STRUCTURE_IN_ANOTHER_ORGANISM
Cytokine response
GOBP_POSITIVE_REGULATION_OF_B_CELL_PROLIFERATION
GOBP_B_CELL_PROLIFERATION
GOBP_REGULATION_OF_PRODUCTION_OF_MOLECULAR_MEDIATOR_OF_IMMUNE_RESPONSE
GOBP_POSITIVE_REGULATION_OF_CELL_POPULATION_PROLIFERATION
GOBP_POSITIVE_REGULATION_OF_PEPTIDYL_TYROSINE_PHOSPHORYLATION
GOBP_TYROSINE_PHOSPHORYLATION_OF_STAT_PROTEIN
GOBP_CELL_SURFACE_RECEPTOR_SIGNALING_PATHWAY_VIA_STAT
GOBP_RESPONSE_TO_CYTOKINE
GOBP_CYTOKINE_MEDIATED_SIGNALING_PATHWAY
GOBP_POSITIVE_REGULATION_OF_CYTOKINE_PRODUCTION
GOBP_CYTOKINE_PRODUCTION
TCR signaling
GOBP_T_CELL_ACTIVATION
GOBP_LYMPHOCYTE_ACTIVATION
GOBP_ADAPTIVE_IMMUNE_RESPONSE
GOBP_CELL_CELL_ADHESION
GOBP_LEUKOCYTE_MEDIATED_IMMUNITY
GOBP_LYMPHOCYTE_MEDIATED_IMMUNITY
GOBP_IMMUNE_EFFECTOR_PROCESS
GOBP_ANTIGEN_RECEPTOR_MEDIATED_SIGNALING_PATHWAY
GOBP_IMMUNE_RESPONSE_REGULATING_CELL_SURFACE_RECEPTOR_SIGNALING_PATHWAY
GOBP_ACTIVATION_OF_IMMUNE_RESPONSE
GOBP_IMMUNE_RESPONSE_REGULATING_SIGNALING_PATHWAY
GOBP_T_CELL_RECEPTOR_SIGNALING_PATHWAY
Killing activity
GOBP_T_CELL_MEDIATED_CYTOTOXICITY
GOBP_T_CELL_MEDIATED_IMMUNITY
GOBP_POSITIVE_REGULATION_OF_CELL_KILLING
GOBP_REGULATION_OF_ALPHA_BETA_T_CELL_ACTIVATION
GOBP_POSITIVE_REGULATION_OF_ALPHA_BETA_T_CELL_ACTIVATION
GOBP_CELL_KILLING
GOBP_LEUKOCYTE_MEDIATED_CYTOTOXICITY
Migration
GOBP_GRANULOCYTE_CHEMOTAXIS
GOBP_GRANULOCYTE_MIGRATION
GOBP_MYELOID_LEUKOCYTE_MIGRATION
GOBP_LEUKOCYTE_CHEMOTAXIS
GOBP_CELL_CHEMOTAXIS
GOBP_LEUKOCYTE_MIGRATION
GOBP_NEUTROPHIL_CHEMOTAXIS
GOBP_NEUTROPHIL_MIGRATION
GOBP_ANTIMICROBIAL_HUMORAL_IMMUNE_RESPONSE_MEDIATED_BY_ANTIMICROBIAL_PEPTIDE
GOBP_ANTIMICROBIAL_HUMORAL_RESPONSE
GOBP_EOSINOPHIL_MIGRATION
GOBP_REGULATION_OF_LYMPHOCYTE_CHEMOTAXIS
−2 −1 0 1 2
NES
GeneRatio
0.2
0.3
0.4
0.5
0.6
.adjust
0.05
0.04
0.03
0.02
0.01
0.00
4 hr
CD8 T killer
CD4 T killer
DP T killer
Cytokine Response
GOBP_POSITIVE_REGULATION_OF_CELL_POPULATION_PROLIFERATION
GOBP_POSITIVE_REGULATION_OF_CYTOKINE_PRODUCTION
GOBP_PEPTIDYL_AMINO_ACID_MODIFICATION
GOBP_B_CELL_PROLIFERATION
GOBP_REGULATION_OF_EPITHELIAL_CELL_DIFFERENTIATION
GOBP_ERK1_AND_ERK2_CASCADE
GOBP_RESPONSE_TO_CYTOKINE
GOBP_CYTOKINE_MEDIATED_SIGNALING_PATHWAY
GOBP_EPITHELIUM_DEVELOPMENT
GOBP_EPITHELIAL_CELL_DIFFERENTIATION
GOBP_PRODUCTION_OF_MOLECULAR_MEDIATOR_OF_IMMUNE_RESPONSE
GOBP_REGULATION_OF_PRODUCTION_OF_MOLECULAR_MEDIATOR_OF_IMMUNE_RESPONSE
GOBP_RESPONSE_TO_WOUNDING
GOBP_RESPONSE_TO_INTERLEUKIN_15
Inflammation
GOBP_DISRUPTION_OF_ANATOMICAL_STRUCTURE_IN_ANOTHER_ORGANISM
GOBP_REGULATION_OF_LYMPHOCYTE_CHEMOTAXIS
GOBP_CENTRAL_NERVOUS_SYSTEM_NEURON_DIFFERENTIATION
Lipid Metabolism
GOBP_CELLULAR_LIPID_METABOLIC_PROCESS
GOBP_ACETYL_COA_BIOSYNTHETIC_PROCESS
DNA Synthesis
GOBP_MEIOSIS_I_CELL_CYCLE_PROCESS
GOBP_INTERSTRAND_CROSS_LINK_REPAIR
GOBP_CHROMOSOME_SEGREGATION
GOBP_DNA_TEMPLATED_DNA_REPLICATION
Migration
GOBP_GRANULOCYTE_MIGRATION
GOBP_NEUTROPHIL_MIGRATION
GOBP_LEUKOCYTE_CHEMOTAXIS
Mitosis
GOBP_MITOTIC_SPINDLE_ASSEMBLY
GOBP_SPINDLE_ASSEMBLY
GOBP_MITOTIC_SISTER_CHROMATID_SEGREGATION
GOBP_MITOTIC_NUCLEAR_DIVISION
ATP Synthesis
GOBP_AEROBIC_RESPIRATION
GOBP_OXIDATIVE_PHOSPHORYLATION
GOBP_CELLULAR_RESPIRATION
GOBP_ATP_SYNTHESIS_COUPLED_ELECTRON_TRANSPORT
Killing Activity
GOBP_LEUKOCYTE_MEDIATED_CYTOTOXICITY
GOBP_CELL_KILLING
GOBP_T_CELL_MEDIATED_IMMUNITY
GOBP_T_CELL_MEDIATED_CYTOTOXICITY
GOBP_LYMPHOCYTE_MEDIATED_IMMUNITY
GOBP_ADAPTIVE_IMMUNE_RESPONSE
GOBP_LYMPHOCYTE_ACTIVATION
GOBP_T_CELL_ACTIVATION
TCR signaling
GOBP_ANTIGEN_RECEPTOR_MEDIATED_SIGNALING_PATHWAY
GOBP_T_CELL_RECEPTOR_SIGNALING_PATHWAY
GOBP_IMMUNE_RESPONSE_REGULATING_CELL_SURFACE_RECEPTOR_SIGNALING_PATHWAY
GOBP_ACTIVATION_OF_IMMUNE_RESPONSE
GOBP_IMMUNE_RESPONSE_REGULATING_SIGNALING_PATHWAY
Phosphorylation
GOBP_PEPTIDYL_TYROSINE_MODIFICATION
GOBP_REGULATION_OF_PEPTIDYL_TYROSINE_PHOSPHORYLATION
GOBP_POSITIVE_REGULATION_OF_PEPTIDYL_TYROSINE_PHOSPHORYLATION
GOBP_CELL_SURFACE_RECEPTOR_SIGNALING_PATHWAY_VIA_STAT
GOBP_TYROSINE_PHOSPHORYLATION_OF_STAT_PROTEIN
GOBP_POSITIVE_REGULATION_OF_PROTEIN_MODIFICATION_PROCESS
GOBP_REGULATION_OF_PHOSPHORYLATION
GOBP_POSITIVE_REGULATION_OF_PHOSPHORUS_METABOLIC_PROCESS
GOBP_POSITIVE_REGULATION_OF_PHOSPHORYLATION
−2 −1 0 1 2
NES
GeneRatio
0.2
0.3
0.4
0.5
0.6
p.adjust
0.05
0.04
0.03
0.02
0.01
0.00

**Supplementary Figure 12. GO biological processes enriched in CD8$^+$, CD4$^+$ or DP killers**

Mutual gene expression difference was calculated using FindAllMarkers() with the settings of logfc.threshold =0 and min.pct =0.20. Bonferroni-adjusted *P* value (two-sided) were used for GSEA as described in Methods. To combine GO terms of high similarity, GO terms aligned to the same phylo tree branch were re-annotated as a single term.

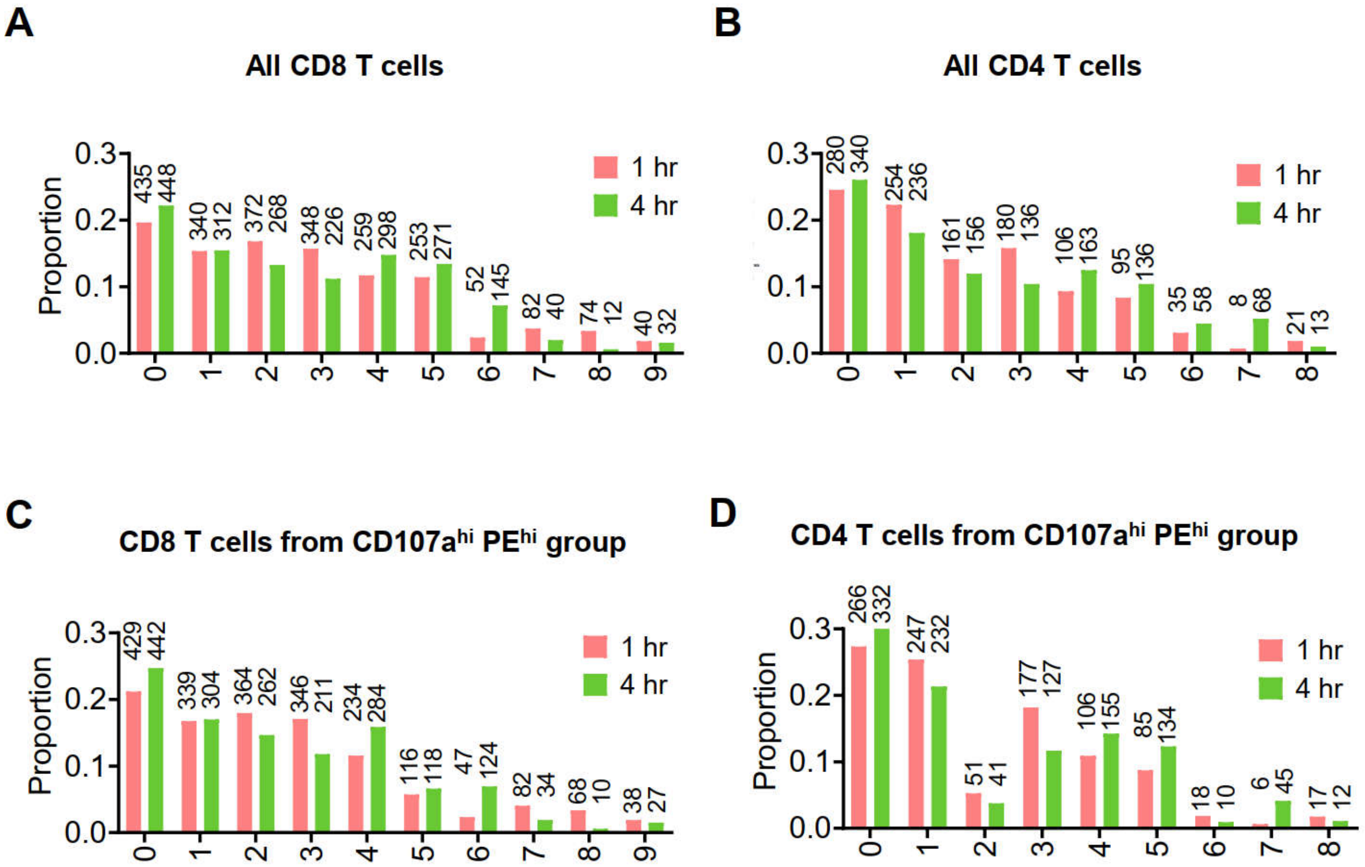


**Supplementary Figure 13. Composition of T-cell subclusters in different time points samples**

(**A**-**B**) Total $CD8^+$ or $CD4^+$ CAR-T cells from both 1 hr and 4 hr were integrated by gene expression as described in Methods, and shown were the proportion and absolute cell counts of identified T cell subclusters in each time point. (**C**-**D**) Likewise, the proportion and absolute cell count of identified T cell subclusters among $CD107a^{hi}$ $PE^{hi}$ CAR-T population in each individual time point.

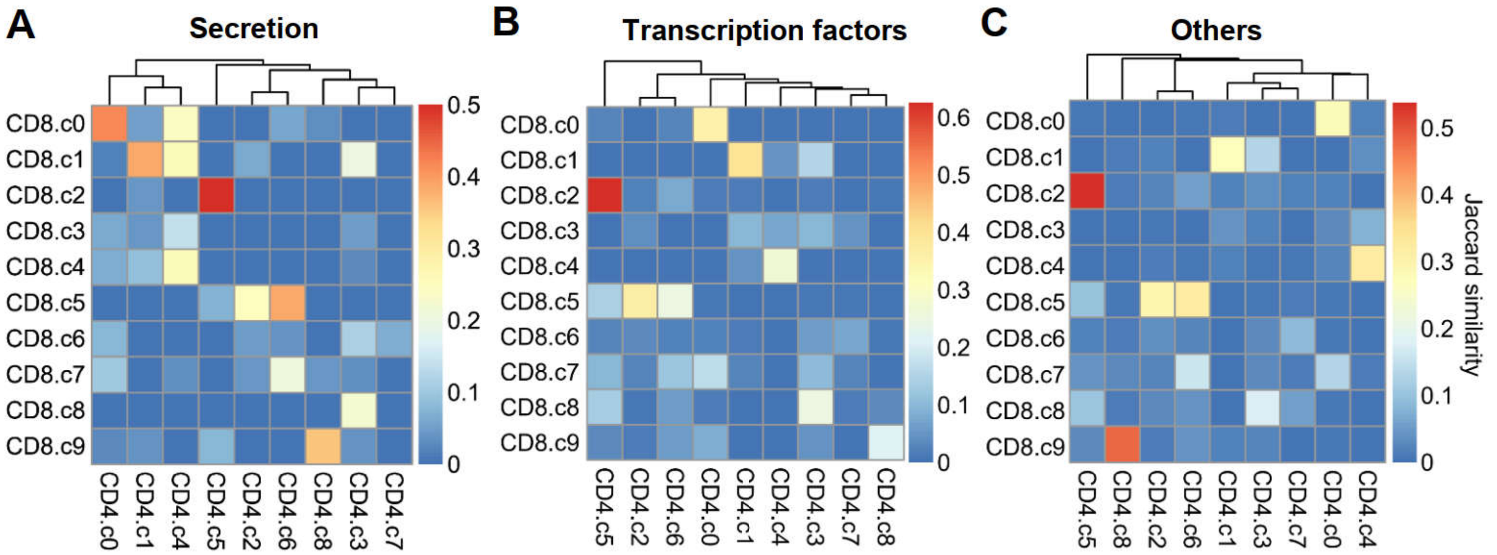


**Supplementary Figure 14. Jaccard similarity of differentially expressed genes between CD4⁺ and CD8⁺ T-cell subclusters**

Wilcoxon rank-sum test using FindAllMarkers() was performed to find differentially expressed genes for the identified $CD8^+$ or $CD4^+$ T cell subclusters separately, with the setting of min.pct =0.20 and logfc.threshold =0.5. Genes passing Bonferroni-adjusted significance test ($P_{adj}$ <0.05) were grouped into secretion-encoding genes (**A**), TF-encoding genes (**B**), and all remaining genes (**C**). Pairwise Jaccard similarity between $CD8^+$ and $CD4^+$ T cell clusters was calculated for each grouped gene signature.

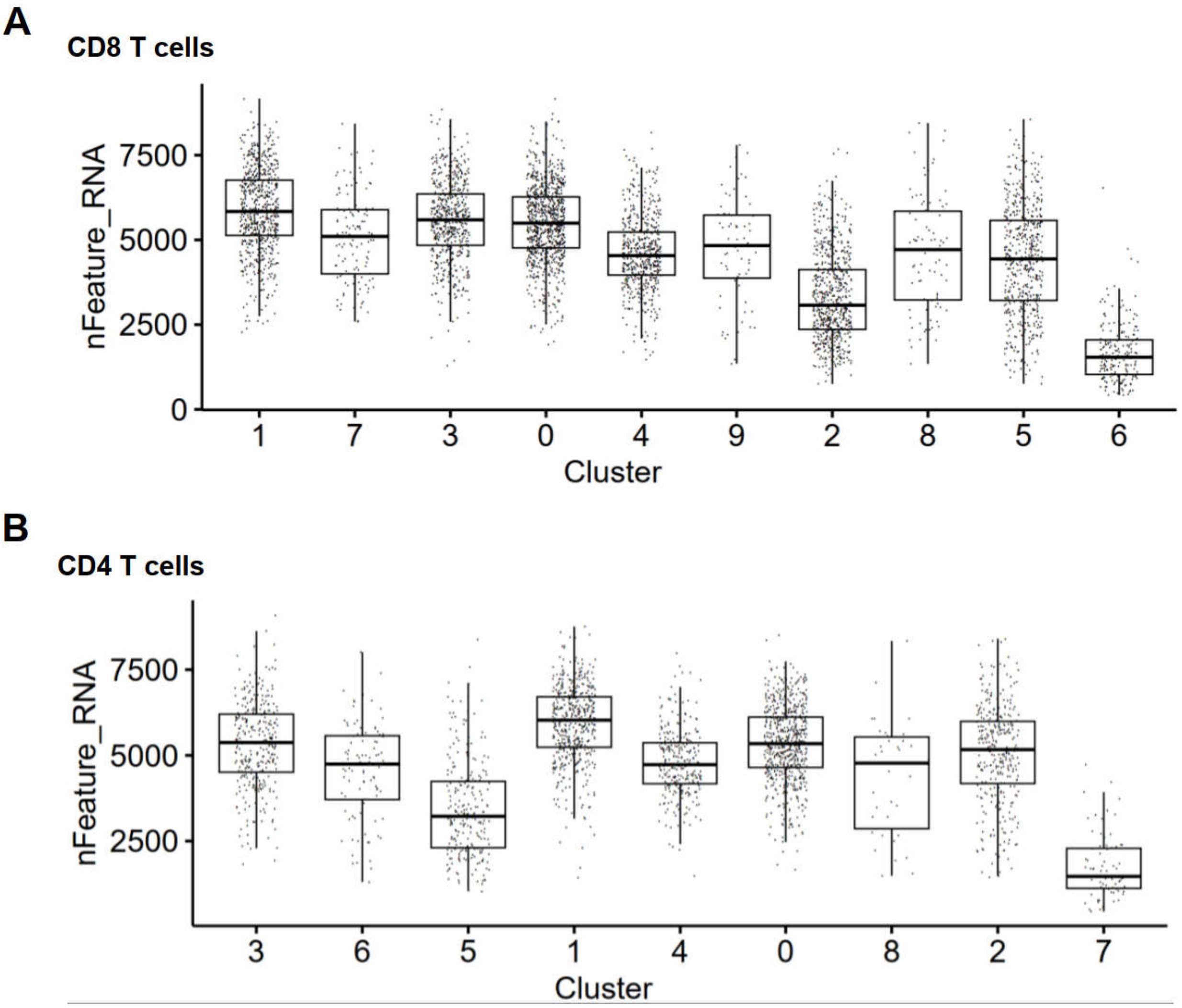


**Supplementary Figure 15. The absolute gene numbers detected among CD4⁺ or CD8⁺ T-cell subclusters**

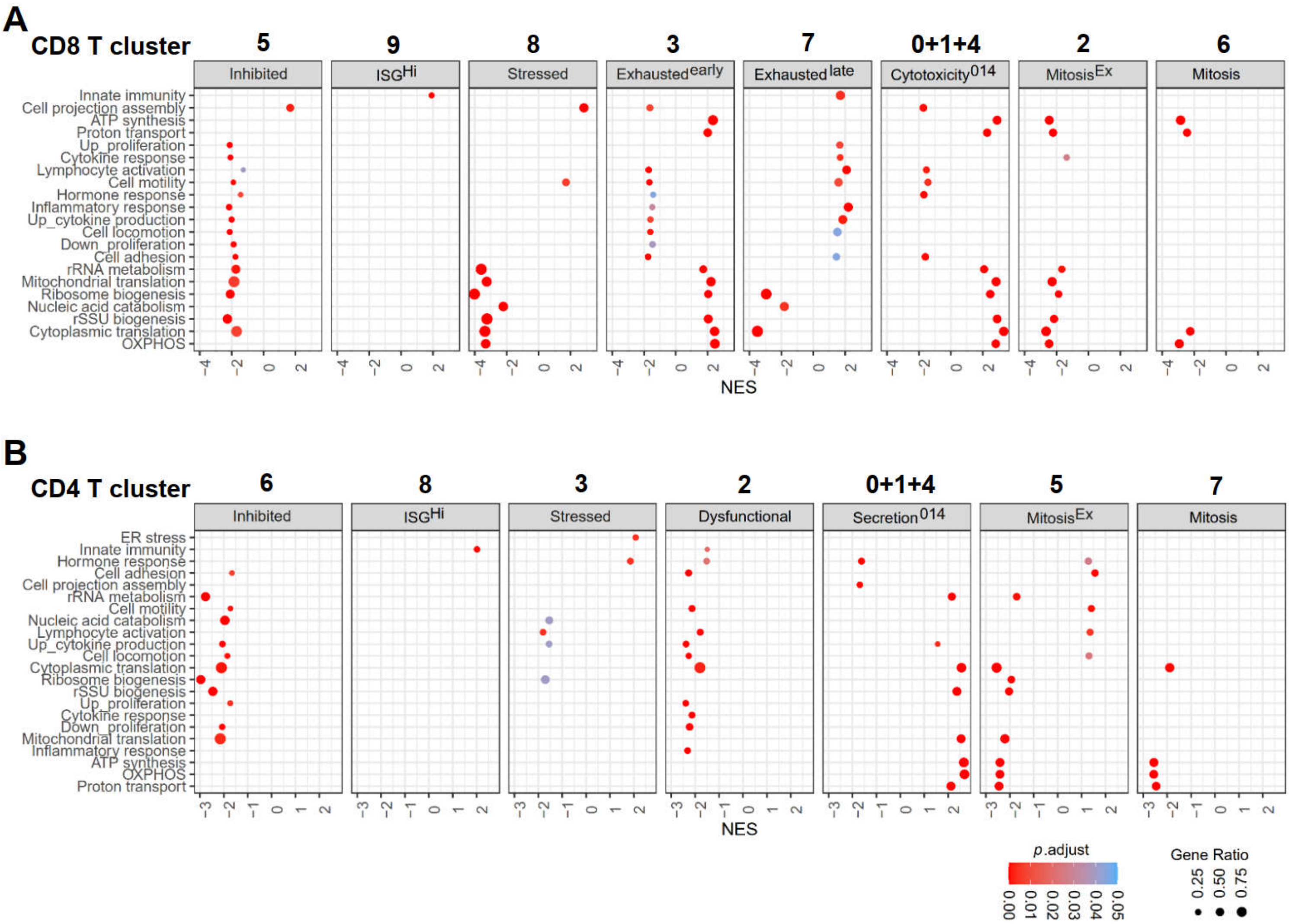


**Supplementary Figure 16. GO biological processes enriched in $CD8^+$ or $CD4^+$ CAR-T subclusters**

Mutual gene expression difference was calculated using FindAllMarkers() with the settings of logfc.threshold =0 and min.pct =0.20. Bonferroni-adjusted *P* value (two-sided) were used for GSEA as described in Methods. The most potent $CD8^+$ cytotoxic CAR-T cells (including $Cytotoxicity^{I}$, $Cytotoxicity^{II}$, $Cytotoxicity^{0}$) or $CD4^+$ cytotoxic CAR-T cells (including $Secretion^{Bal}$, $Secretion^{I}$, $Secretion^{I}$) were combined into one single group.

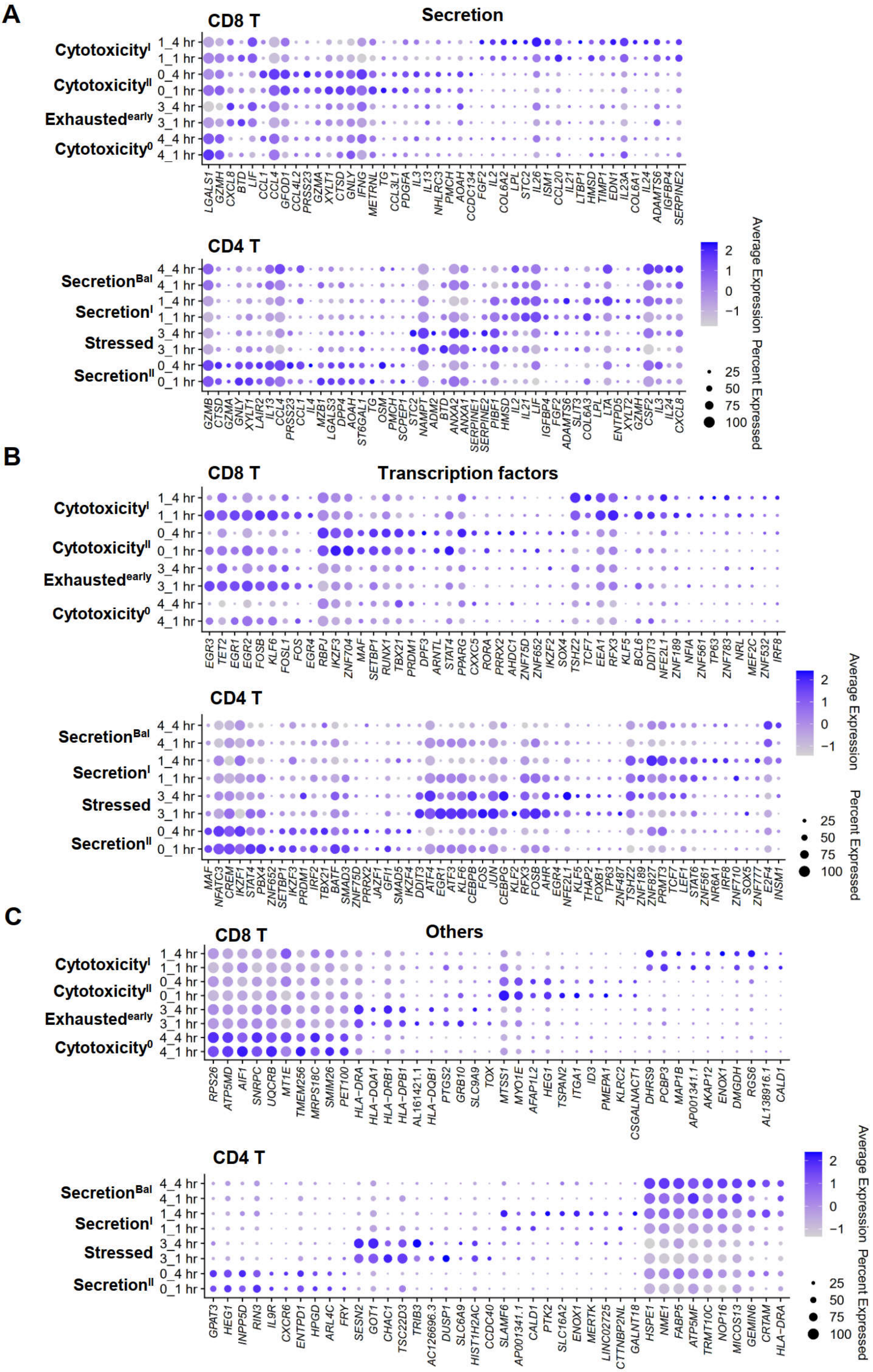
A
Secretion
CD8 T
Cytotoxicity^I
Cytotoxicity^II
Exhausted^early
Cytotoxicity^0
1_4 hr
1_1 hr
0_4 hr
0_1 hr
3_4 hr
3_1 hr
4_4 hr
4_1 hr
LGALS1 GZMH CXCL8 BTD LIF CCL1 CCL4 GFOD1 CCL4L2 PRSS23 GZMA XYLT1 CTSD GNLY IFNG METRNL TG CCL3L1 PDGFA IL3 IL13 NHLRC3 PMCH AOAH CCDC134 FGF2 IL2 COL6A2 LPL STC2 IL26 ISM1 CCL20 IL21 LTBP1 HMSD TIMP1 EDN1 IL23A COL6A1 IL24 ADAMTS6 IGFBP4 SERPINE2
CD4 T
Secretion^Bal
Secretion^I
Stressed
Secretion^II
GZMB CTSD GZMA GNLY XYLT1 LAIR2 IL13 CCL4 PRSS23 CCL1 IL4 MZB1 LGALS3 DPP4 AOAH ST6GAL1 TG OSM PMCH SCPEP1 STC2 NAMPT ADM2 BTD ANXA2 ANXA1 SERPINE1 SERPINE2 PIBF1 HMSD IL2 IL21 LIF IGFBP4 FGF2 ADAMTS6 SLIT3 COL6A3 LPL LTA ENTPD5 XYLT2 GZMH CSF2 IL3 IL24 CXCL8
Average Expression
2
1
0
−1
Percent Expressed
25
50
75
100
B
Transcription factors
CD8 T
EGR3 TET2 EGR1 EGR2 FOSB KLF6 FOSL1 FOS EGR4 RBPJ IKZF3 ZNF704 MAF SETBP1 RUNX1 TBX21 PRDM1 DPF3 ARNTL STAT4 PPARG CXXC5 RORA PRRX2 AHDC1 ZNF75D ZNF652 IKZF2 SOX4 TSHZ2 TCF7 EEA1 RFX3 KLF5 BCL6 DDIT3 NFE2L1 ZNF189 NFIA ZNF561 TP63 ZNF783 NRL MEF2C ZNF532 IRF8
CD4 T
MAF NFATC3 CREM IKZF1 STAT4 PBX4 ZNF652 SETBP1 IKZF3 PRDM1 IRF2 TBX21 BATF SMAD3 ZNF75D PRRX2 JAZF1 GFI1 SMAD5 IKZF4 DDIT3 ATF4 EGR1 ATF3 KLF6 CEBPB FOS JUN CEBPG KLF2 RFX3 FOSB AHR EGR4 NFE2L1 KLF5 THAP2 FOXB1 TP63 ZNF487 TSHZ2 ZNF189 ZNF827 PRMT3 TCF7 LEF1 STAT6 ZNF561 NR6A1 IRF8 ZNF710 SOX5 ZNF777 E2F4 INSM1
C
Others
CD8 T
RPS26 ATP5MD AIF1 SNRPC UQCRB MT1E TMEM256 MRPS18C SMIM26 PET100 HLA−DRA HLA−DQA1 HLA−DRB1 HLA−DPB1 AL161421.1 HLA−DQB1 PTGS2 GRB10 SLC9A9 TOX MTSS1 MYO1E AFAP1L2 HEG1 TSPAN2 ITGA1 ID3 PMEPA1 KLRC2 CSGALNACT1 DHRS9 PCBP3 MAP1B AP001341.1 AKAP12 ENOX1 DMGDH RGS6 AL138916.1 CALD1
CD4 T
GPAT3 HEG1 INPP5D RIN3 IL9R CXCR6 ENTPD1 HPGD ARL4C FRY SESN2 GOT1 CHAC1 TSC22D3 TRIB3 AC126696.3 DUSP1 SLC6A9 HIST1H2AC CCDC40 SLAMF6 AP001341.1 CALD1 PTK2 SLC16A2 ENOX1 MERTK LINC02725 CTTNBP2NL GALNT18 HSPE1 NME1 FABP5 ATP5MF TRMT10C NOP16 MICOS13 GEMIN6 CRTAM HLA−DRA

**Supplementary Figure 17. Gene expression signatures of the most potent killer cell clusters**

For the most potent $CD8^+$ cytotoxic CAR-T cells clusters(Cytotoxicity$^{I}$, Cytotoxicity$^{II}$, Cytotoxicity$^{0}$, and Exhausted$^{early}$) or $CD4^+$ cytotoxic CAR-T cells clusters(Secretion$^{Bal}$, Secretion$^{I}$, Secretion$^{II}$, and Stressed), mutual gene expression difference was calculated using FindAllMarkers() with the settings of logfc.threshold =0.5 and min.pct =0.20. Genes passing Bonferroni-adjusted significance test ($P_{adj}$ <0.05) were grouped into secretion-encoding genes (**A**), TF-encoding genes (**B**), and all remaining genes(**C**). Shown were the expression of top genes for each classification.

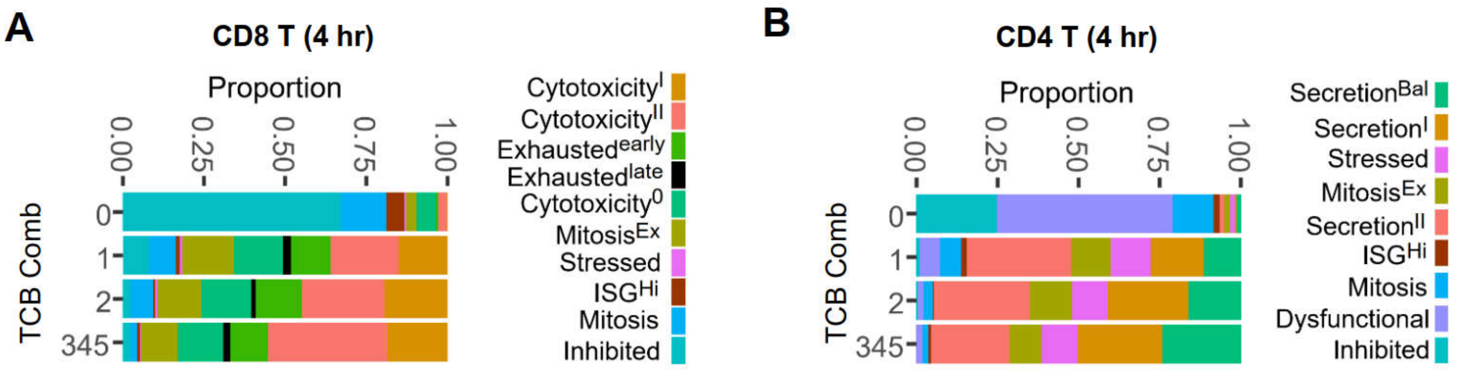


**Supplementary Figure 18. Cell composition among different combination of TCB capture**

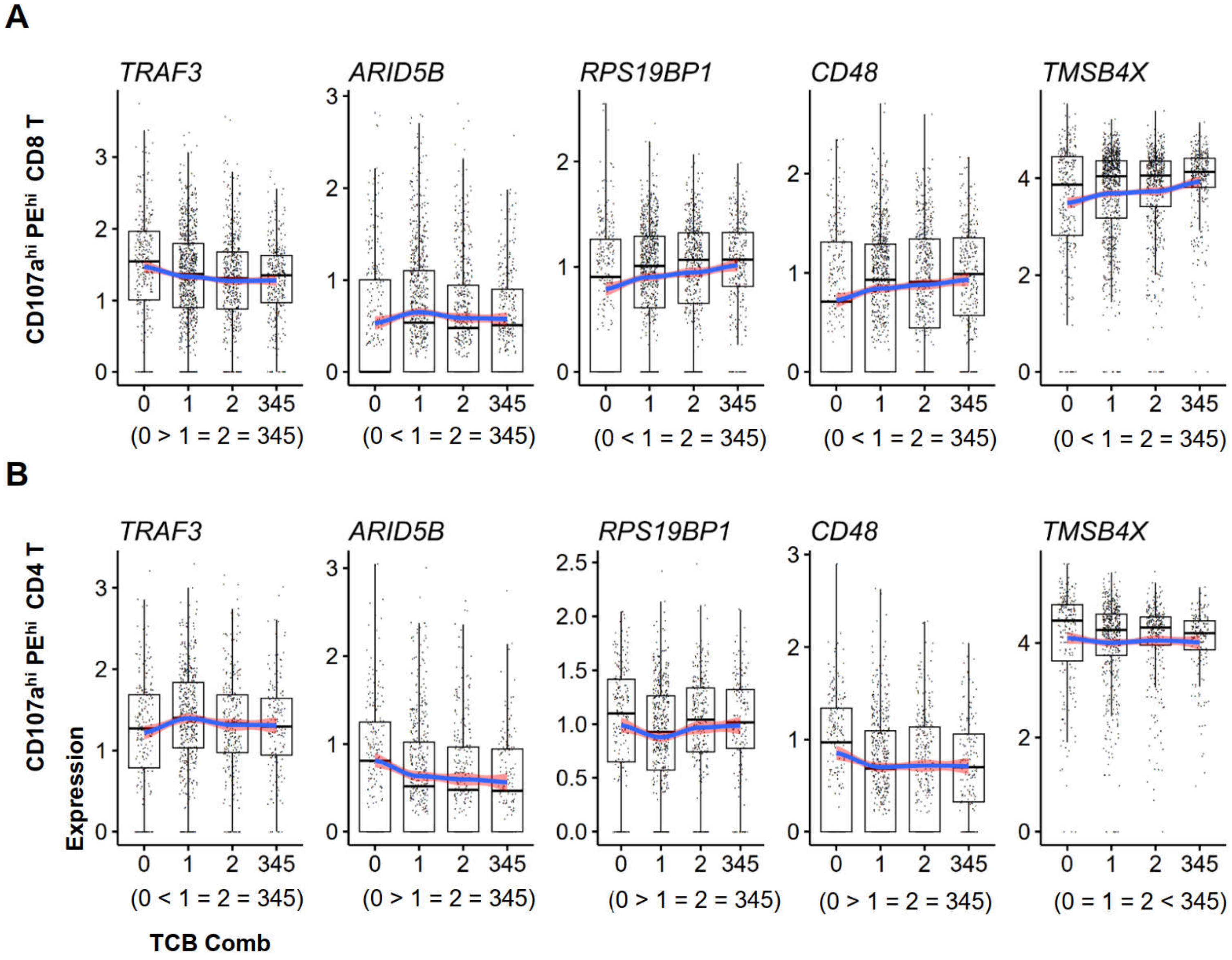


**Supplementary Figure 19. Genes of opposite expression dynamics between CD8⁺ and CD4⁺ CAR-T cells during serial killing acquisition**

The significance of differential gene expression between two adjacent groups was tested using Wilcoxon method with Holm correction. The symbol “>”, “<”, and “=” indicated the trend of gene expression according to significance test ($P_{adj}$ <0.05).
